# C5aR1 signaling promotes region and age dependent synaptic pruning in models of Alzheimer’s Disease

**DOI:** 10.1101/2023.09.29.560234

**Authors:** Angela Gomez-Arboledas, Maria I. Fonseca, Enikö Kramar, Shu-Hui Chu, Nicole Schartz, Purnika Selvan, Marcelo A. Wood, Andrea J. Tenner

## Abstract

**INTRODUCTION:** Synaptic loss is a hallmark of Alzheimer’s disease (AD) that correlates with cognitive decline in AD patients. Complement-mediated synaptic pruning has been associated with this excessive loss of synapses in AD. Here, we investigated the effect of C5aR1 inhibition on microglial and astroglial synaptic pruning in two mouse models of AD.

**METHODS:** A combination of super-resolution and confocal and tridimensional image reconstruction was used to assess the effect of genetic ablation or pharmacological inhibition of C5aR1 on the Arctic48 and Tg2576 models of AD.

**RESULTS:** Genetic ablation or pharmacological inhibition of C5aR1 rescues the excessive pre-synaptic pruning and synaptic loss in an age and region dependent fashion in two mouse models of AD, which correlates with improved long-term potentiation (LTP).

**DISCUSSION:** Reduction of excessive synaptic pruning is an additional beneficial outcome of the suppression of C5a-C5aR1 signaling, further supporting its potential as an effective targeted therapy to treat AD.

## 1. Background

Alzheimer’s Disease (AD) is a devastating neurodegenerative disorder that impairs memory and results in cognitive deficits, often accompanied by psychiatric disorders. Neuropathologically, AD is characterized by the extracellular accumulation of amyloid-ß in senile plaques and neurofibrillary tangles and synaptic and neuronal loss (1). It is also well established that inflammation and the complement system play a key role in disease progression, where glial cell responses to amyloid may contribute to neuronal damage and cognitive impairment (2–4). In fact, GWAS studies have identified at least three complement genes as being significantly associated with AD, CLU, CR1 and C1S (5). Activation of the classical and alternative complement pathways during Alzheimer’s disease occurs as a response to fibrillar ß-sheet amyloid ß (fAß) and hyperphosphorylated tau (6–8). Additionally, multiple complement mediators have been found co-localizing with Aß plaques, not only in mouse models (9, 10) of the disease but more importantly, in human post-mortem samples from AD patients (11, 12), further supporting a critical role of the complement system in AD pathogenesis (reviewed in (13)). Activation of the complement system by fibrillar Aß could ultimately generate C5a which can then interact with a proinflammatory C5aR1 receptor (which is upregulated by injury or infection) (14, 15) on microglial cells, triggering a potent inflammatory response.

During development, synapse elimination or refinement is a necessary process to ensure the appropriate establishment of synaptic circuits. The classical pathway of the complement system, through C1q and C3, has been directly linked to beneficial synaptic pruning that occurs in developmental stages, including adulthood (16, 17). However, multiple studies implicate synaptic loss as a hallmark of Alzheimer’s disease that correlates best with the cognitive decline in AD patients (18). Interestingly, complement-mediated synaptic pruning has been associated with the excessive synaptic loss that occurs during Alzheimer’s disease. It has been hypothesized that excessive complement activation could mediate aberrant synapse pruning by microglial engulfment in AD models, as well as in other neurodegenerative disorders (19–23), ultimately leading to cognitive decline.

It is known that C1q binds to phosphatidyl serine and other damage exposed molecules on apoptotic cells, and likely interacts with similar molecules at weak or nonactive synapses (24–26). Activation of complement at the synapses can lead to microglial engulfment of C1q/iC3b tagged synapses via CR3 receptor on microglial cells (20). In addition, deficiency or blockage of C1q, C3, C4 and CR3 has been shown to suppress synaptic pruning and prevent cognitive deficits in several mouse models of AD (19, 21, 27, 28). Furthermore, evidence for a neuroprotective role for C1q suggests that upstream inhibition of the complement system might not be an optimal target to treat AD (29, 30).

Previous studies in our lab and others, have shown that the genetic or pharmacological inhibition of C5a-C5aR1 signaling reduced pathology, and rescued synaptic and cognitive loss in several mouse models of AD (31–35), suggesting that C5a-C5aR1 signaling might be a better therapeutic target than blocking the upstream components of the complement cascade. Moreover, two different C5aR1 antagonist have already been proven safe in humans in different clinical trials for autoimmune disorders, further emphasizing the potential for C5aR1 as a therapeutic target (36, 37). In addition, it is well established that astrocytes play a key role in circuit refinement during development and adulthood (38), however, their role in the excessive synaptic loss observed in AD is still controversial. In a recent report, astrocytes were found to preferentially ingest excitatory synapses while microglia engulfed inhibitory synapses in mouse models of tauopathy and AD, (39), suggesting that astrocytes might also have an important role in synaptic pruning during Alzheimer’s disease.

Therefore, here we investigated the effect of genetic ablation or pharmacological inhibition of C5aR1 on microglial and astroglial synaptic pruning in two different mouse models of Alzheimer’s disease. We show that synaptic loss and excessive microglial synaptic pruning can be partially rescued by genetic deletion or pharmacological inhibition of C5aR1 in an age and region dependent manner.

## 2. Methods

### 2.1 Animals

All animal experimental procedures were approved by the Institutional Animal Care and Use Committee of University of California, Irvine, and performed in accordance with the NIH Guide for the Care and Use of Laboratory Animals. The AD mouse model Arctic48 (Arc), initially provided by Dr. Lennart Mucke (Gladstone Institute, San Francisco, CA, USA), contains the human APP transgene with the Indiana (V717F), Swedish (K670 N + M671 L), and Arctic (E22G) mutations (under the control of the platelet derived growth factor-ß promoter), and thus produces Aß protofibrils and fibrils at 2–4 months old (Cheng et al, 2004). The C5aR1 knockout mice, generated by target deletion of the C5a receptor gene and initially provided by Dr. Rick Wetsel (University of Texas, Houston), were crossed with Arctic^+/−^ mice, kindly provided by Dr. Lennart Mucke (Gladstone Institute), to produce Arctic mice lacking C5aR1 (Arc-C5aR1KO) and wild type littermate mice lacking the C5a receptor (C5aR1KO) (**Supp Figure 1A**). Tg2576 mice, developed by K. Hsiao (40) overexpressed the 695 isoform of the amyloid precursor protein (APP) with the Swedish mutation (KM670/671NL) under the control of the prion promoter on a B6/SJL genetic background. Only female hemizygous Tg2576 mice were used as they develop cortical amyloid plaques by 11-13 months of age, earlier than male Tg2576 (40). WT (B6/SJL) female littermates were used as control mice (**Supp Figure 1B**).

### 2.2 C5aR1 antagonist (PMX205) treatment

C5aR1 antagonist, PMX205, was kindly provided by Dr. Ian Campbell, Teva Pharmaceuticals, West Chester, PA. Treatment with PMX205 was performed as previously described (34). Briefly, PMX205 was administered in the drinking water at 20 μg/ml to mice at the onset of amyloid pathology at 12 months of age for the Tg2576 mice (and B6/SJL WT littermates) (**Supp Figure 1B**). PMX205 treatment was administered for 12 weeks and mice were singly housed and had free access to the drinking water during the whole treatment.

### 2.3 Tissue collection and preparation

WT, C5aR1KO, Arc and Arc-C5aR1KO mice at different ages (2.7, 5, 7 and 10 months-old) were deeply anesthetized with isoflurane and transcardially perfused with 4% paraformaldehyde/PBS and post fixed overnight in 4% paraformaldehyde/PBS at 4^°^C or perfused with PBS and fixed for 24hs with 4% paraformaldehyde/PBS at 4^°^C (no differences were seen in results from different perfusion methods). Brains were sectioned at 40 μm using a vibratome (Leica VT 1000S). For the Tg2576 and WT littermates that underwent PMX205 treatment, mice were deeply anesthetized with isoflurane and transcardially perfused with HBSS modified buffer (Hank’s Balanced Salt Solution without Calcium and Magnesium), containing Actinomycin D (5 μg/ml) and Triptolide (10 μM). Half brains were fixed in 4% paraformaldehyde for 24h and sectioned at 40 μm thickness in the coronal plane on a vibratome (Leica VT1000S).

### 2.4 Golgi staining and Sholl analysis

Mice were perfused as above, and half brains were fixed and, after sectioning (150 μm), stained using the super-Golgi Kit (Bioenno Lifesciences, Santa Ana, CA), following the instructions of the manufacturer. All sections were coded, such that the analysis was done blinded. The stained tissues were then quantified using a stereology Zeiss Axio Imager M2 with NeuroLucida software Version 11.03. The soma and apical dendrites along with its dendritic branching were traced under 100× magnification in the CA3 region of the hippocampus for each animal followed by Sholl analyses, determining the average number of branching for a given distance from the soma (μm) of each neuron. The average number of branchings for a given distance from the soma for the neurons of each animal and genotype +/− SEM were generated using GraphPad Prism.

### 2.5 Antibodies

The following primary antibodies were used for this study: mouse monoclonal anti synaptophysin (SP) (clone SVP38, 1.4 ug/ml, Sigma), guinea pig polyclonal anti VGlut11 (1:1000, Millipore), rabbit polyclonal anti postsynaptic density 95 (PSD95, 1 ug/ml, Invitrogen), rat monoclonal anti CD68 (clone FA-11, 1.4 ug/ml, Bio-Rad), rabbit polyclonal anti Iba-1 (1:1000, Wako), rat monoclonal anti C3 (clone 11H9, 0.5ug/ml, Abcam) and rabbit monoclonal anti C1q (clone 27.1, hybridoma supernatant previously described (41, 42)). The specificity of the anti-mouse C1q antibody was demonstrated by the lack of C1q staining detected in the hippocampal CA1-SR of an Arctic C1q conditional KO (ArcC1qa^FL/FL^:Cx3cr1^ERT2^) compared with the Arc C1qa^FL/FL^ at super-resolution level (**Supp Figure 2**) and characterized previously (41).

### 2.6 Immunofluorescence

Coronal brain sections were blocked with 2% BSA, 10% normal goat serum, 0.1% Triton, TBS (1h, RT), and incubated with primary antibodies diluted in blocking solution overnight at 4^°^C. After washing, antigen-bound primary antibodies were detected with the corresponding labelled Alexa 488, Alexa 555 or Alexa 647 secondary antibodies (1/300 – 1/500, Invitrogen) that were incubated for 1h at RT. Sections were then mounted either with ProLong glass antifade mounting media (Invitrogen) for super-resolution microscopy or with Vectashield Hardset (Vector) for confocal imaging.

### 2.7 Image acquisition

For super-resolution imaging, hippocampal regions CA1-SR, DG-ML, and CA3-SL of WT, C5aR1-KO, Arc and Arc-C5aR1KO at different ages were acquired with a Zeiss LSM 880 Airyscan microscope and Zen image acquisition software (Zeiss) with identical conditions. Images for each animal and within all the regions of interest were collected using a 63x oil objective and 3-4 Z stacks (187 nm step interval, within a depth of 3 μm, covering an area of 58×58 μm) were obtained. For super-resolution imaging of WT and Tg2576 mice treated with/without PMX205, hippocampal regions CA1-SR and CA3-SL were acquired by Super-Resolution Lattice Structured Illumination Microscopy (Lattice-SIM) using an Elyra 7 microscope system (Zeiss). Images for each animal and within all the regions of interest were collected using a 63x 1.4NA Plan-Apo objective lens and Immersol 518F immersion oil. 4 z-stacks (110 nm step interval, within a depth of 5-8 μm, covering an area of 64×64 μm) per mouse/group were acquired. Images were processed using ZEN SIM^2^ on the ZEN black edition software. For confocal imaging of microglial/astroglial synaptic pruning, individual microglial/astroglial cell images were obtained with a Leica TCS SP8 microscope under a 63x objective and a 3.5 zoom factor. Z-stacks were acquired in 0.3 µm steps to image the whole microglial/astroglial cell. For each mouse the CA1-SR and CA3-SL regions of the hippocampus were imaged. A total of 15 cells per mouse, per region and age were acquired and analyzed.

### 2.8 Imaris quantitative analysis

Image analyses were carried out in several hippocampal regions by using Imaris 9.2 or 9.5 software (Bitplane Inc). Colocalization of pre and post synaptic puncta (SP or VGlut1 with PSD95) and C1q or C3 associated with presynaptic puncta (SP or VGlut1) were quantified using the spots or the surfaces function on Imaris and Matlab software to determine the total number of colocalized spots (defined at ≤ 200 nm distance) or the number of spots close to surfaces (≤ 200 nm distance). Spots were normalized to the total image volume. Synaptic engulfment was defined as the co-localization (defined at ≤ 200 nm distance) of VGlut1+ synapses (detected by Imaris spots) with either CD68 (%VGlut1 engulfment in microglial/astroglial lysosomes) or Iba1/GFAP (%VGlut1 engulfment) surfaces and normalized to the total volume of the image.

### 2.9 Hippocampal slice preparation and LTP recording

Hippocampal slices were prepared from male and female 10-month-old WT, Arctic, WT-C5aR1KO and Arctic-C5aR1KO mice (n =34). Following isoflurane anesthesia, mice were decapitated, and the brain was quickly removed and submerged in ice-cold, oxygenated dissection medium containing (in mM): 124 NaCl, 3 KCl, 1.25 KH_2_PO_4_, 5 MgSO_4_, 26 NaHCO_3_, and 10 glucose. Coronal hippocampal slices (340 µm) were prepared using a vibratome (VT1000S) before being transferred to an interface recording containing preheated artificial cerebrospinal fluid (aCSF) of the following composition (in mM): 124 NaCl, 3 KCl, 1.25 KH_2_PO_4_, 1.5 MgSO_4_, 2.5 CaCl_2_, 26 NaHCO_3_, and 10 glucose and maintained at 31 ± 1^0^C. Slices were continuously perfused with this solution at a rate of 1.75-2 ml/min while the surface of the slices were exposed to warm, humidified 95% O_2_ / 5% CO_2_. Recordings began following at least 2 hr of incubation.

Field excitatory postsynaptic potentials (fEPSPs) were recorded from CA1b stratum radiatum using a single glass pipette filled with 2M NaCl (2-3 MΩ) in response to orthodromic stimulation (twisted nichrome wire, 65 µm diameter) of Schaffer collateral-commissural projections in CA1 stratum radiatum. Pulses were administered at 0.05 Hz using a current that elicited a 50% maximal response. Paired-pulse facilitation was measured at 40, 100, and 200 sec intervals prior to setting baseline. After establishing a 20 min stable baseline, long-term potentiation (LTP) was induced by delivering 5 ‘theta’ bursts, with each burst consisting of four pulses at 100 Hz and the bursts themselves separated by 200 msec (i.e., theta burst stimulation or TBS). The stimulation intensity was not increased during TBS. Data were collected and digitized by NAC 2.0 Neurodata Acquisition System (Theta Burst Corp., Irvine, CA) and stored on a disk.

### 2.10 Statistics

All data were analyzed using either a t-test for comparison of two groups and one-way ANOVA or two-way ANOVA, followed by Tukey’s post hoc test for comparison among more than 2 groups by using GraphPad Prism Version 9 (La Jolla, CA) and Microsoft Excel. The significance was set at 95% confidence. *p<0.05, **p<0.01, ***p<0.001 and ****p<0.0001. Data were presented as the mean ± SEM (standard error of the mean).

## 3. Results

### 3.1 C5aR1 depletion on the Arctic model of AD restores short- and long-term plasticity at 10 months of age

Genetic ablation of C5aR1 in the Arctic mouse model of AD results in a protection from cognitive decline and from loss of neuronal complexity in the CA1 (31) and CA3 region of the hippocampus **(Supp Fig 3**) at 10 months of age. To evaluate the potential contributions of C5aR1 in synaptic plasticity, the cellular mechanism believed by many to underlie memory processes, we first examined long-term potentiation (LTP) in acute hippocampal slices from 10 months WT, C5aR1KO, Arctic and Arctic-C5aR1KO mice. Evoked field excitatory post-synaptic potentials (fEPSPs) were recorded from the CA1b stratum radiatum hippocampal region. Following a train of theta burst stimulation (TBS) to a collection of Schaffer-commissural fibers, fEPSP slope increased immediately and dramatically in WT slices that then decayed over the following 10 min to reach a stable level of potentiation at approximately 50% above baseline 60 min post-induction (**Fig. 1A-C**). However, LTP in slices from C5aR1KO mice was significantly reduced 50-60 min post-TBS compared to WT slices (**Fig. 1A-C**). Importantly, LTP in slices from Arctic-C5aR1KO were indistinguishable from WT controls. Accordingly, a marked deficit in LTP was found in slices from Arctic mice (**Fig. 1A-C**). These results indicate that like the C5aR1KO mouse, the Arctic mouse model has a strong impairment in LTP, but more importantly, the deletion of C5aR1KO in the Arctic mouse model of AD can restore deficits in LTP to control levels.

**Figure 1:**
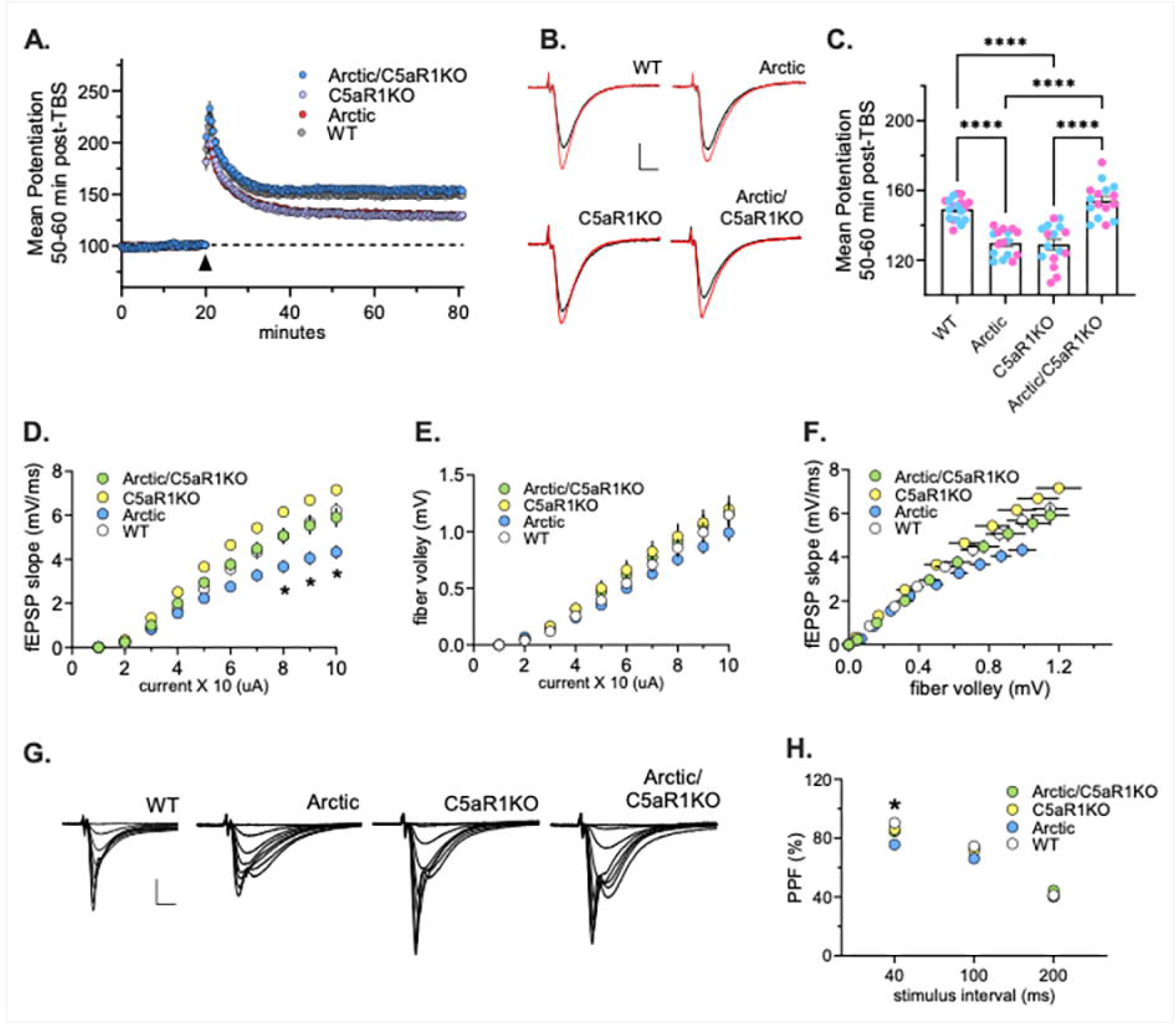
Genetic deletion of C5aR1 rescues LTP deficit in Arctic mice at 10 months of age. **A.** Hippocampal slices were collected from 10-month-old (mo) male and female WT, Arctic, C5aR1KO and Arctic/C5aR1KO mice. Long-term potentiation (LTP) in the CA1-SR area was measured. Following a 20 min stable recording, LTP was induced by applying TBS (black arrow) and recording of baseline stimulation was resumed for an additional 60 min. Immediately following TBS, fEPSP slope increased and then slowly decayed to a stable level of LTP above baseline recordings in slices from WT (white circle), Arctic (grey circle), C5aR1KO (light blue), and Arctic/C5aR1KO (dark blue) mice. **B.** Representative traces collected during baseline (black line) and 60 min post-TBS (red line). Scale: 1 mV/5 ms. **C.** The mean potentiation 50-60 min post-TBS was significantly impaired in Arctic and C5aR1KO mice relative to WT and Arctic/C5aR1KO mice (p<0.0001). Light blue and pink circles represent mean potentiation of each slice in male and female mice, respectively. **D-E.** Input/output curves assessed the amplitude of the fEPSP slope (panel D) and fiber volley (panel E) across a range of stimulus currents (10-100 uA) (*p<0.01). **F.** Input/output curves comparing the amplitudes of the presynaptic fiber volley to the fEPSP slope across a range of stimulus currents. **G.** Representative field recordings collected during the first through eight steps of the input/output curve in slices from WT, Arctic, C5aR1KO and Arctic/C5aR1KO mice. Scale: 2 mV/5 ms. **H.** Paired pulse facilitation (PPF) of the initial slope of the synaptic response was compared in slices from Arctic/C5aR1KO and C5aR1KO relative to WT mice, and in Arctic mice at the 40 ms stimulus interval (p = 0.03).

We then investigated if the absence of C5aR1 affects any of the conventional measures of transmission. Input-output curves measuring the magnitude of the fEPSP slope (**Fig. 1D-G**) and fiber volley (**Fig 1E-G**) in response to an increase in stimulus current were comparable for WT, C5aR1KO and Artic-C5aR1KO mice, but there was a marked depression of fEPSP slope in Arctic mice (**Fig 1D**). These results indicate that deletion of C5aR1 alone does not affect axon excitability. However, we did find that slices from the Arctic mice (relative to controls) showed depressed axon excitability and that deletion of C5aR1KO from the Arctic mouse restores normal axonal transmission. We then tested for differences in transmitter release using paired pulse facilitation (PPF). Slices from C5aR1KO and Arctic/C5aR1KO did not differ from WT controls (**Fig. 1H**), but PPF in Arctic slices was significantly reduced at 40 ms stimulus interval suggesting that Arctic mice have a reduction in transmitter mobilization. Taken all together, our results showed that while the Arctic mice showed the greatest deficit in both synaptic transmission and LTP, combining this transgene with C5aR1KO appears to have restored both short- and long-term plasticity.

### 3.2 Absence of C5aR1 rescues the excessive VGlut1 presynaptic puncta loss at 10 months of age in the Arctic mouse model of AD

Here we assessed whether genetic ablation of C5aR1 would similarly dampen synaptic pruning by microglia at 10 months of age in the CA3-SL hippocampal region. Super-resolution microscopy (**Fig 2A1**) and Imaris quantification (**Fig 2A2-A4**) of VGlut1, PSD95 and VGlut1-PSD95 colocalized synaptic puncta showed a significant VGlut1 presynaptic loss (55% reduction) in the Arctic mouse model of AD (vs WT). Moreover, we observed an increased in VGlut1+ synaptic puncta in the Arc-C5aR1KO mice when compared to Arctic mice, indicative of a partial presynaptic puncta rescue when C5aR1 was depleted (**Fig 2A2**). No differences on any of the genotypes were observed on PSD95 postsynaptic density (**Fig 2A3**). Colocalization of VGlut1 and PSD95 synaptic puncta showed a significant synaptic loss (40% reduction) in the AD mice when compared to WT (**Fig 2A4**), however, no rescue of synaptic density was detectable. These results indicate a potential role of C5a-C5aR1 signaling in the excessive presynaptic loss associated with Alzheimer’s disease (43).

**Figure 2:**
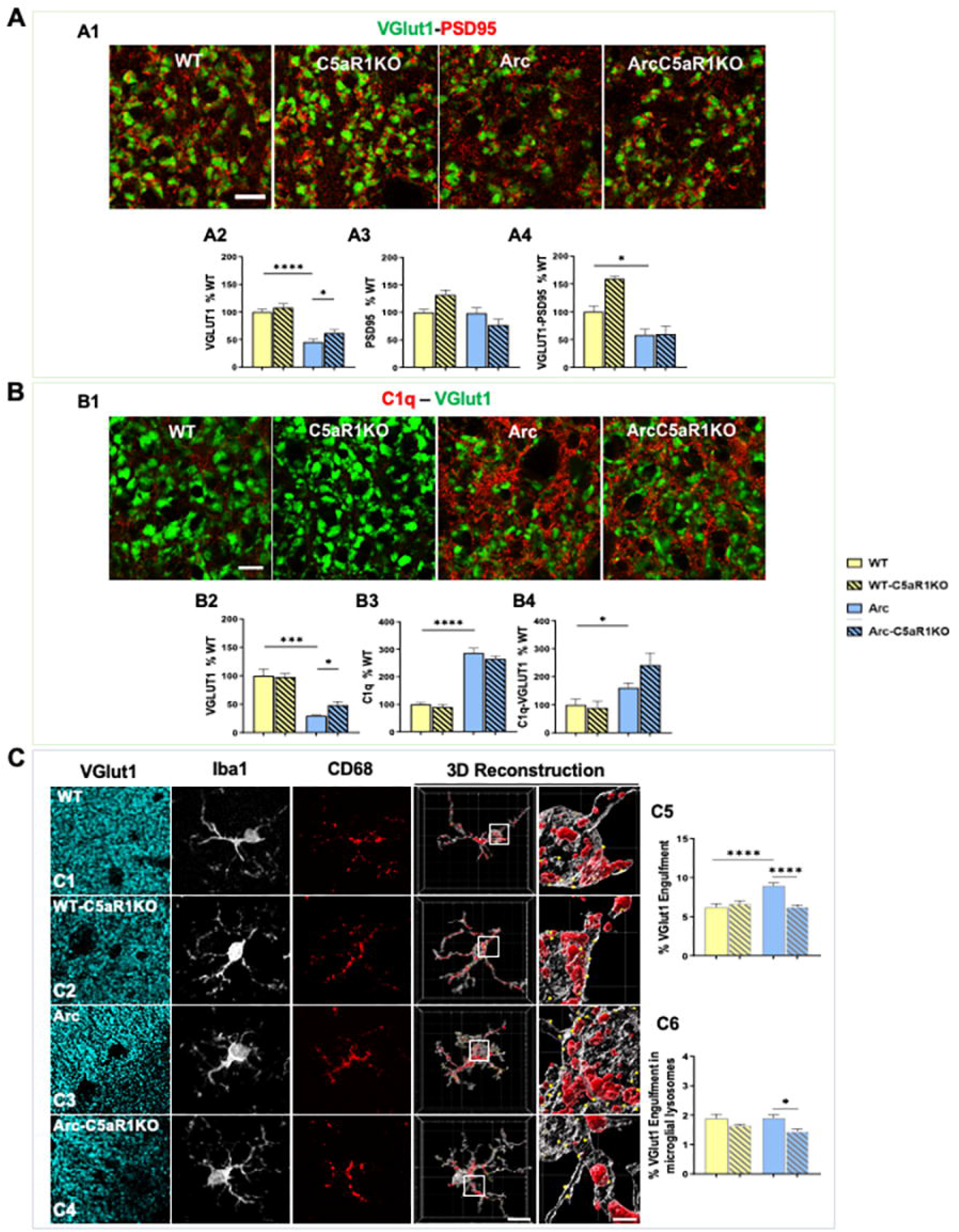
Lack of C5aR1 rescues VGlut1 presynaptic loss and reduces microglial engulfment despite higher C1q tagging in the CA3-SL hippocampal region at 10 months of age. **A.** Representative super-resolution images of VGlut1 (green) and PSD95 (red) synaptic puncta in the CA3-SL of 10m Arc, Arc-C5aR1KO and their respective WT littermates. Scale bar: 5 μm (A1). Imaris quantitative analysis of VGlut1, PSD95 and colocalized synaptic puncta (A2-A4). Data are shown as Mean ± SEM (normalized to WT control group) of 3-4 images per animal and 2-6 animals per genotype. **B**. Representative super-resolution images of C1q (red) and VGlut1 (green) synaptic puncta. Scale bar: 5 μm (B1). Imaris quantification of C1q, VGlut1 and C1q-VGlut1 colocalization (B2-B4). Data are shown as Mean ± SEM (normalized to WT control group) of 3 images per animal and 4-5 animals per genotype. **C.** Confocal images and 3D reconstruction with surface rendering using Imaris software of individual microglial cells (Iba1+; grey), presynapses (VGlut1+; cyan), and lysosomes (CD68+; red). Scale bar: 10 μm; inserts 2 μm (C1-C4). Quantitative analysis of VGlut1+ presynaptic puncta engulfment per microglia (C5). Quantitative analysis of VGlut1+ presynaptic puncta localized within the microglial lysosomes (C6). Data are shown as Mean ± SEM of 15 individual microglial cells/mouse and n=3 mice per genotype. *p<0.05, ** p<0.01, *** p<0.001, **** p<0.0001.

It has been widely described that weak or less-active synapses associated with complement components C1q and cleaved C3 are actively engulfed by microglial cells through CR3, a receptor for the opsonic complement activation product iC3b. To determine if C1q tagging correlated with the changes observed in synaptic density, and if the presence of C1q could be used to predict vulnerable or reduced synaptic puncta, we analyzed VGlut1 and C1q immunostaining (**Fig 2B1**) in WT, C5aR1KO, Arc and Arc-C5aR1KO mice at 10 months of age in the CA3-SL region. Quantitative analysis from super-resolution images showed a significant increase (2.7-2.9-fold increase) in C1q in Arctic mice relative to WT (**Fig 2B3**). When assessing the colocalization of C1q with VGlut1, we observed a significant increase in C1q-VGlut1 colocalized puncta in the Arctic mice (vs WT), which was even further increased in the Arc-C5aR1KO (although not statistically significant when compared to Arc mice) (**Fig 2B4**). The lack of correlation of our results here, where VGlut1 puncta is increased in Arc-C5aR1KO (compared to Arctic mice) but there is no difference in % C1q puncta, suggest that although C1q tagging plays a role in presynaptic pruning in this AD mouse model, it is not sufficient.

C3 has been shown to be involved in synapse pruning (19–21, 44). To investigate if C3 tagging would correlate with the VGlut1 pre-synaptic density changes observed at 10 months of age in the Arc and Arc-C5aR1KO (vs wildtype) mice in the CA3-SL region, sections were immunostained for C3 and VGlut1, and quantification studies were carried out with Imaris software (**Supp Fig 4**). As seen in previous experiments (**Fig 2A2**), VGlut1 density was decreased in the Arc relative to WT and partially rescued in the Arc-C5aR1KO (**Supp Fig 4A2**). C3 staining, using an antibody that recognizes whole C3 as well as its cleavage products, was observed mainly on astrocytes and deposited on the neuropil. Our results showed a high amount of C3 positive astrocytes, mainly in the vicinity of amyloid plaques (data not shown), which parallels previous bulk RNAseq and IHC results from our lab showing a strong astroglial response (characterized by an increased in GFAP and C3) in the hippocampus of Arctic and Arc-C5aR1KO mice at 7 and 10 months of age (35). Here, our results at 10 months in the CA3-SL region showed a significant increase of C3 puncta (excluding astroglia-associated C3) in the Arc mice when compared to WT littermates, which was further increased in the Arc-C5aR1KO mice (**Supp Fig 4A3**). However, no significant differences were found in the C3-VGlut1 colocalized puncta among any of the different genotypes (**Supp Fig 4A4**), suggesting that as with C1q, while C3 may play a role in synaptic pruning in the Arctic mouse model at 10 months of age, other factors must also be contributing substantially to the ingestion of presynaptic VGlut1.

Since complement activation at the synapses or activation by fibrillar amyloid plaques will generate C5a (when C5 is present) and since the expression of C5aR1 increases with age and pathology (14), to determine if the absence of C5aR1 results in a reduced microglial phagocytosis of the synapses, we co-stained VGlut1 with the microglial markers Iba1 and CD68 (**Fig 2 C1-C4**). Single microglial cells (and thus not plaque associated) in the CA3-SL region from all four genotypes were imaged and co-localization of presynaptic markers within the microglial cell body or within the microglial lysosomes were quantified using Imaris. Our results showed an increased amount of VGlut1 puncta within the microglial cell body in the Arctic mice (vs WT), consistent with an increase in microglial phagocytosis of synapses. However, the absence of C5aR1 in the Arc mouse model restored the VGlut1 engulfment down to WT levels (**Fig 2C5**), which correlated with the rescue of VGlut1 presynaptic density observed in the Arc-C5aR1KO mice (vs Arctic) discussed above (**Fig 2A2**), suggesting that the excessive microglial engulfment observed in the Arctic mouse model of AD could be the main driver of the presynaptic loss described. No changes between WT and Arc mice were observed when the amount of VGlut1 puncta in the microglial lysosomes was analyzed, although a significant decrease was observed when comparing Arc-C5aR1KO vs Arctic mice (**Fig 2C6**), suggesting that microglial cells in the Arc-mice might be more efficient in the degradation of the engulfed material in response to the severe amyloid pathology described in this AD model. Together, our results demonstrate a role of C5a-C5aR1 signaling in the excessive microglial presynaptic pruning that leads to synaptic loss and ultimately cognitive deficit at late stages of the pathology in the Arctic mouse model of Alzheimer’s Disease.

### 3.3 The beneficial effects of C5aR1 genetic ablation on the rescue of VGlut1 excessive presynaptic loss are dependent on the hippocampal region at 10 months of age

Since C5aR1 deletion reduced the loss of neuronal complexity at 10 months of age in both the CA1-SR (31) and CA3-SL (**Supp Fig 3**), we assessed the synaptic density in the CA1-SR region at 10 months of age, similarly to our analysis in the CA3-SL region described above. As expected at 10 months of age, a significant reduction in synaptic density was observed in the CA1-SR hippocampal region of Arc mice (compared to WT littermates) (**Fig 3A4**). However, in contrast to the CA3-SL region, while there was a significant and substantial decrease (41% reduction) in SP presynaptic puncta in Arctic mice (vs WT), it was not rescued by deletion of C5aR1 (Arc-C5aR1KO vs Arc mice) (**Fig 3A2**). As in our previous results in the CA3-SL, we did not observe any differences in the amount of PSD95 postsynaptic puncta among any of the genotypes at 10 months of age in the CA1-SR (**Fig 3A3**). SP-PSD95 colocalized puncta was significantly reduced in both Arc (49% reduction) and Arc-C5aR1KO (45% reduction) mice when compared to their control groups (**Fig 3A4**), suggesting that absence of C5aR1 did not have a significant effect in the CA1-SR hippocampal region. Similar results were observed when quantifying VGlut1 puncta and VGlut1-PSD95 colocalization in the same groups, age, and CA1-SR hippocampal region (data not shown). Similar to our previous results in the CA3-SL region, decreases were observed in both SP and VGlut1 presynaptic puncta in CA1-SR in the Arc and Arc-C5aR1KO mice that were accompanied by a significant increase in C1q deposition (**Fig 3B-C**) at the presynapses.

**Figure 3:**
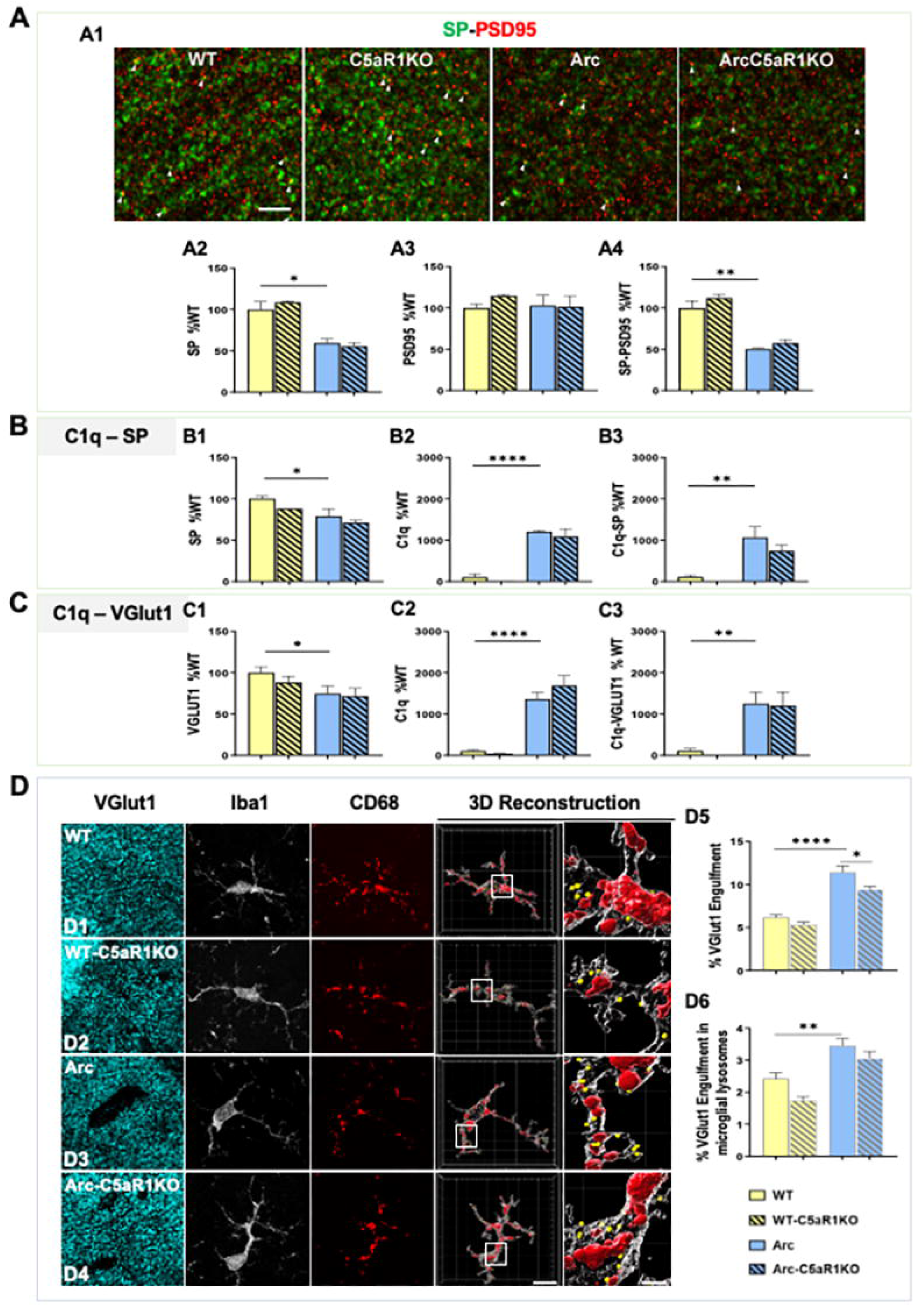
Absence of C5aR1 in the Arctic model of AD fails to rescue presynaptic loss in the CA1-SR hippocampal region at 10 months of age. **A.** Representative super-resolution images of SP (green) and PSD95 (red) at 10 months of age in CA1-SR (arrowheads show colocalization of pre and post synaptic puncta) Scale bar: 5 μm. (A1). Quantitative analysis of SP, PSD95 and colocalized SP-PSD95 puncta (A2-A4). Data are shown as Mean ± SEM (normalized to WT control group) of 4-6 images per animal and 2-4 animals per genotype. **B**. Imaris quantification of SP, C1q, and C1q-SP colocalization from super-resolution images in CA1-SR. Data are shown as Mean ± SEM (normalized to WT control group) of 3 images per animal and 1-4 mice per genotype. **C**. Imaris quantification of VGlut1, C1q, and C1q-VGlut1 colocalization from super-resolution images in CA1-SR. Data are shown as Mean ± SEM (normalized to WT control group) of 3 images per animal and 3-7 mice per genotype. **D**. Confocal images and 3D reconstruction with surface rendering using Imaris of individual microglial cells (Iba1+; grey), presynapses (VGlut1+; cyan), and lysosomes (CD68+; red). Scale bar: 10 μm; inserts 2 μm. (D1-D4). Quantitative analysis of VGlut1+ presynaptic puncta engulfment per microglia (D5). Quantitative analysis of VGlut1+ presynaptic puncta localized within the microglial lysosomes (D6). Data are shown as Mean ± SEM of 15 individual microglial cells/mouse and n=3 mice per genotype. *p<0.05, ** p<0.01, *** p<0.001, **** p<0.0001.

Double labelling of C3 and VGlut1 in the CA1-SR hippocampal region showed that while there were decreases in total SP or VGlut1 puncta in the Arc mice as seen in **Fig 3B or 3C**, no significant differences in the total amount of C3 or C3 colocalized with VGlut1 were seen in any of the genotypes analyzed (**Supp Fig 4B)**. Interestingly, in a different hippocampal region (DG-ML), despite the substantial increases in C1q and C1q tagged VGlut1 and SP synapses (**Supp Fig 5B)**, no changes in synaptic puncta were observed across the different genotypes (**Supp Fig 5A**), further suggesting that changes in synaptic puncta are region specific and that C1q or iC3b tagging alone is not sufficient to trigger synaptic engulfment and synaptic loss.

Analysis of VGlut1+ synapses engulfed by microglial cells in the CA1-SR region showed a significant reduction of VGlut1 puncta inside microglial cells in the absence of C5aR1 in the Arctic model (Arc-C5aR1KO), when compared to C5aR1 sufficient Arctic mice (**Fig 3D5**). These results suggest that even if microglial synaptic pruning is reduced by C5aR1 deletion, this beneficial reduction is not enough to rescue the excessive presynaptic loss observed at 10 months of age in the Arctic model of AD in the CA1-SR region of the hippocampus.

Overall, our results demonstrated that C5a-C5aR1 signaling affects microglial synaptic engulfment and presynaptic loss in the Arctic mouse model of Alzheimer’s Disease in a region dependent manner.

### 3.4 C5a-C5aR1 influences synaptic loss and microglial synaptic engulfment in a region-specific dependent manner at 7 months of age

While behavioral deficits in the Arctic model were only observed at 10 months of age (31), we investigated if synaptic loss and microglial synaptic pruning might precede cognitive deficit at an earlier stage of the disease. Super-resolution microscopy showed a significant decrease in VGlut1 puncta (55% reduction) (**Fig 4A1-A2**) and colocalized VGlut1-PSD95 puncta (47% reduction) (**Fig 4A4**) in the Arctic mice when compared to WT littermates at 7 months of age in the CA3-SL. Interestingly at this earlier age, Arc-C5aR1KO mice showed a trend for a partial rescue of this synaptic density (**Fig 4A2, A4**), when compared to Arc mice. At this same age, the microglial engulfment of VGlut1 pre-synapses showed a significant increase in the Arctic genotype (vs WT) with a trend for a reduction of VGlut1 puncta detected within the microglial cells in the Arc-C5aR1KO mice (**Fig 4C5**), thus correlating with the trend for an improvement in synaptic density observed in the Arc-C5aR1KO relative to the C5aR1 sufficient Arctic mice (**Fig 4A2 and 4A4**). No differences in the amount of VGlut1 puncta within the microglial lysosomes were found in any of the genotypes, suggesting that at 7 months of age, the lysosomal function is not affected by the amyloid pathology present in the Arc mouse model of AD (**Fig 4C6**). Furthermore, increased levels of C1q associated with VGlut1 presynaptic puncta were observed in the Arctic mice when compared to WT **(Fig 4B**), which correlates with the decrease in VGlut1 (**Fig 4A2**), colocalized VGlut1-PSD95 puncta (**Fig 4A4**) and VGlut1 microglial engulfment (**Fig 4C5**). However, since the amount of C1q puncta is the same in the Arc and ArcC5aR1KO, changes in C1q tagging does not account for the partial rescue of the synaptic density (**Fig. 4A4**) and synaptic engulfment observed in the absence of C5aR1 (Arc-C5aR1KO vs Arc mice).

**Figure 4:**
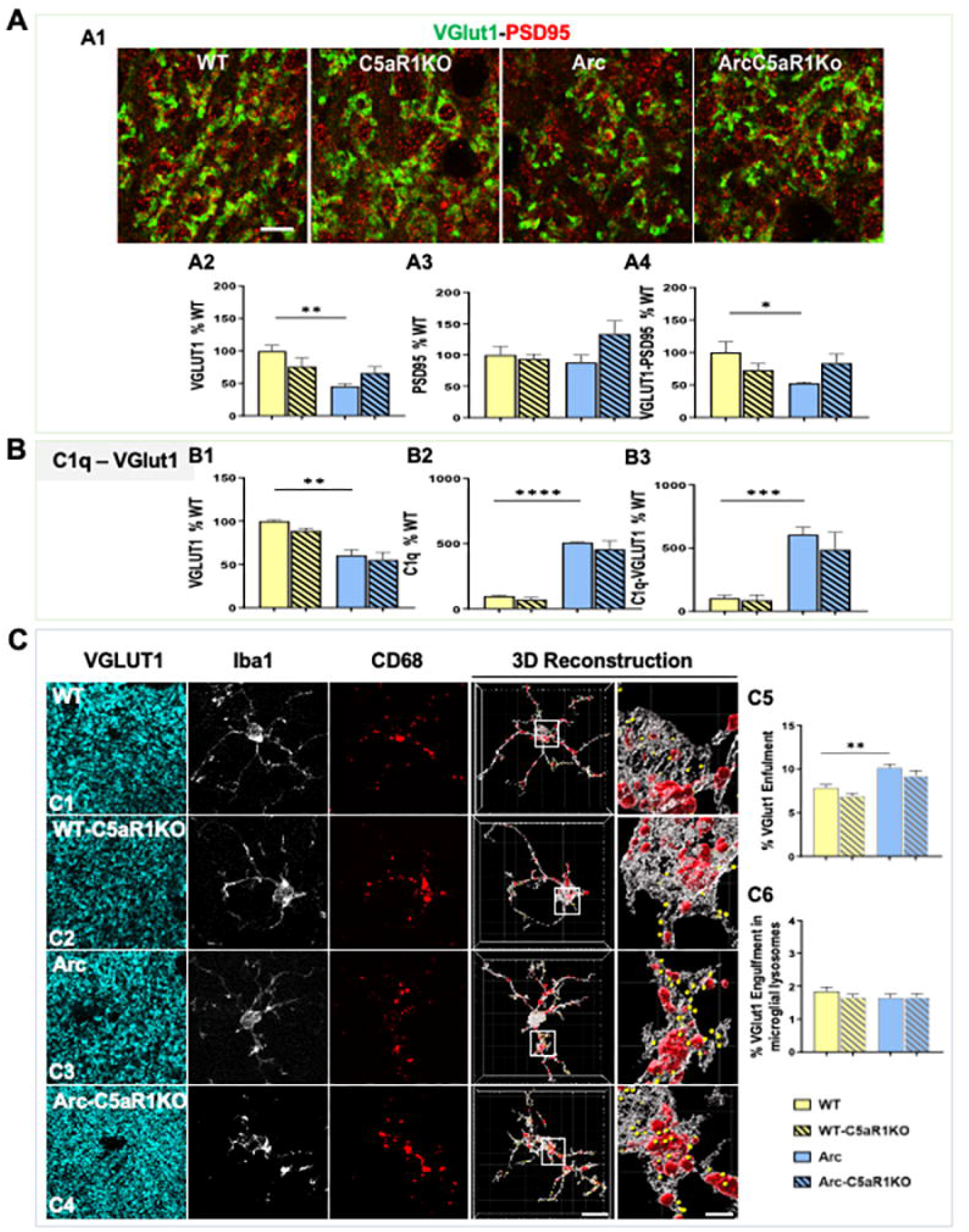
VGlut1 presynaptic loss in the Arctic mouse model of AD is not reversed in the absence of C5aR1 at 7 months in the CA3-SL region of the hippocampus. **A.** Super-resolution representative images (A1) and quantitative analysis of VGlut1, PSD95 and colocalized VGlut1-PSD95 puncta (A2-A4) in CA3-SL of 7m WT, WT-C5aR1KO, Arc, and Arc-C5aR1KO. Scale bar: 5 μm. Data are shown as Mean ± SEM (normalized to WT control group) of 3 images per animal and n=3-4 per genotype. **B**. Quantitative analysis of VGlut1, C1q and C1q-VGlut1 colocalized puncta in CA3-SL at 7 months of age. Data are shown as Mean ± SEM (normalized to WT control group) of 3 images per animal and n=3-4 per genotype. **C**. Confocal images and 3D reconstruction of Iba1, CD68 and VGlut1 in CA3-SL at 7 months of age in the Arctic mouse model of AD. Scale bar: 10 μm; inserts 2 μm (C1-C4). Quantitative analysis of VGlut1+ presynaptic puncta engulfment per microglia (C5). Quantitative analysis of VGlut1+ presynaptic puncta localized within the microglial lysosomes (C6). Data are shown as Mean ± SEM of 15 individual microglial cells/mouse and n=3-4 mice per genotype. *p<0.05, ** p<0.01, *** p<0.001, **** p<0.0001.

In contrast to what we observed at 7 months of age in the CA3-SL hippocampal region, no differences were found in VGlut1 presynaptic puncta (**Supp Fig 6A1**), VGlut1-PSD95 colocalized puncta (**Supp Fig 6A3**) or VGlut1 microglial engulfment (**Supp Fig 6C5**) in any of the genotypes in the CA1-SR region at the same age, further supporting our previous results at 10 months of age where the role of C5a-C5aR1 in synaptic loss and synaptic engulfment is dependent upon the hippocampal region. In addition, we observed a significant increase in C1q tagged VGlut1 puncta (**Supp Fig 6B2**) in Arctic mice (compared to WT) in the CA1-SR hippocampal region at 7 months of age, which correlates with the increases in C1q-VGlut1 puncta that we observed at both 7 and 10 months of age in the CA1 and CA3 areas. However, no differences were found in the total amount of VGlut1 presynaptic puncta engulfed by microglial cells (**Supp Fig 6C5**). These results further support our hypothesis that C1q tagging is not sufficient to drive the excessive hippocampal synaptic pruning and synaptic loss associated with Alzheimer’s Disease.

Finally, we analyzed even earlier stages of the disease (2.7 and 5 months of age) to assess whether synaptic loss could precede the excessive accumulation of amyloid beta plaques observed in the Arctic model of Alzheimer’s Disease. No changes in synaptic density (VGlut1-PSD95 colocalization) were observed among any of the genotypes at 2.7 (**Supp Fig 7A**) or 5 months of age (**Supp Fig 7B**) in any of the regions analyzed, suggesting that synaptic loss events occur at later stages of the disease, when amyloid plaques are dominant in the Arctic model.

### 3.5 Pharmacological inhibition of C5aR1 rescues the excessive synaptic loss in the Tg2576 mouse model of Alzheimer’s Disease

Besides the beneficial role of genetic ablation of C5aR1 in preventing the loss of neurite complexity, presynaptic loss (data reported in this manuscript) and cognitive deficits in the Arctic mouse model of AD (31, 35), pharmacological inhibition of C5a-C5aR1 signaling with PMX205, a potent C5aR1 antagonist, (33, 34) in two different mouse models of Alzheimer’s disease has shown beneficial effects in modulating gene expression and preventing synaptic and cognitive deficits. Therefore, we analyzed synaptic density changes in the Tg2576 mouse model of Alzheimer’s disease upon PMX205 treatment (**Supp Fig 1B**). VGlut1 and PSD95 synaptic puncta were measured at 15 months of age in both the CA3-SL and CA1-SR hippocampal regions (**Fig 5A-B**). Our results in the CA3-SL area, showed a significant decrease (14% decrease) in VGlut1 presynaptic puncta (**Fig 5A2**) and also in VGlut1-PSD95 colocalized puncta (28% decrease) (**Fig 5A4**) in the Tg2576-H_2_O mice (when compared to WT-H_2_O) that was completely rescued when PMX205 was administered to Tg2576 (Tg2576-PMX205). Similar to our results in the Arctic model, no changes were observed in the PSD95 postsynaptic puncta density among any of the different groups (**Fig 5A3**). Interestingly, our previous results in the CA3-SL region showed an increase in the VGlut1 synaptic engulfment by microglial cells in the Tg2576-H_2_O (vs WT-H_2_O) that was significantly reduced upon treatment with PMX205 (34), together suggesting that C5aR1 antagonism reduces microglial synaptic pruning which results in the rescue of the excessive presynaptic loss and synaptic density observed in the Tg2576 mouse model of Alzheimer’s Disease.

**Figure 5:**
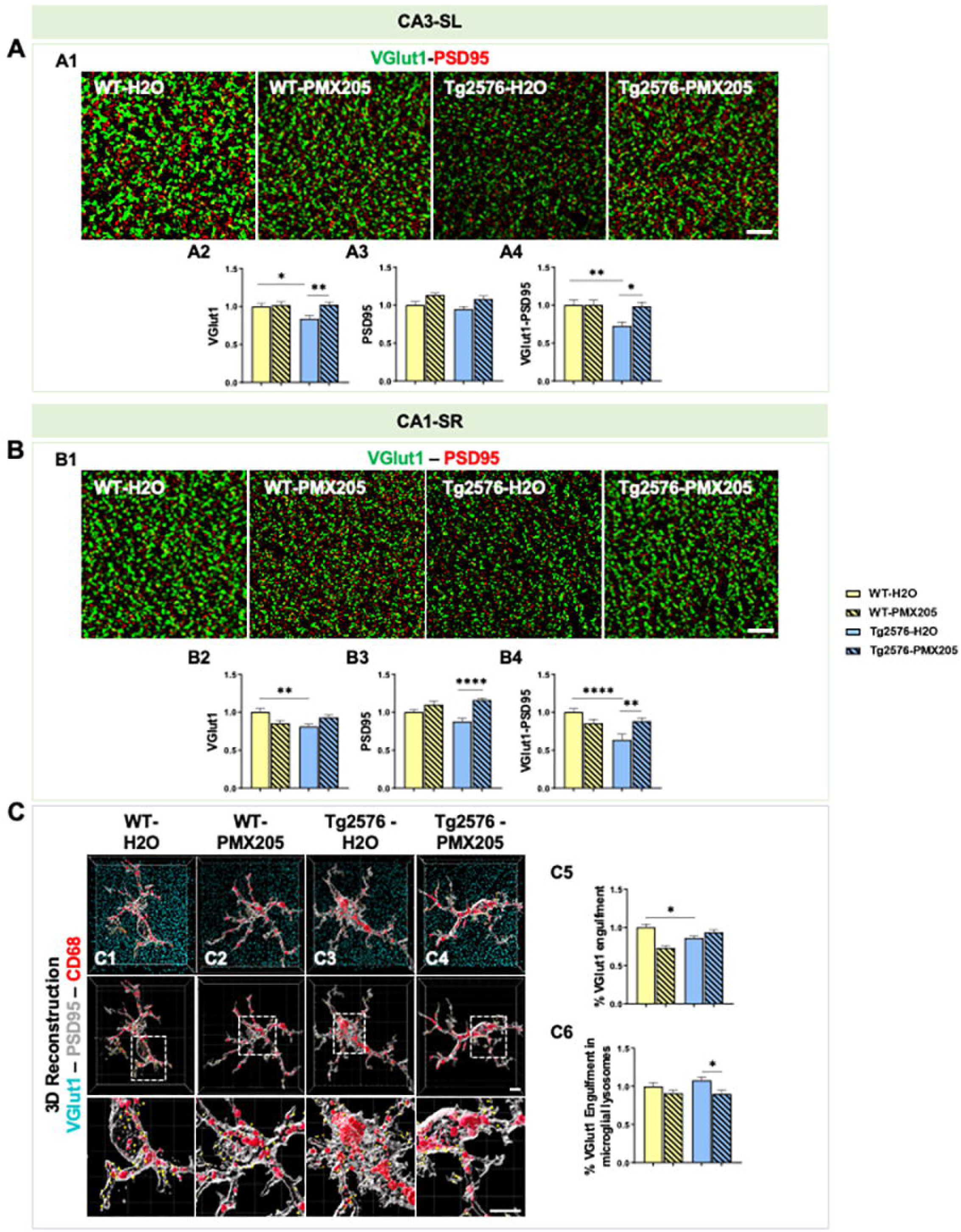
Treatment with PMX205 rescues the excessive synaptic loss observed in the Tg2576 mouse model of Alzheimer’s Disease. **A-B**. Representative super-resolution images of VGlut1 (green) and PSD95 (red) puncta in the CA3-SL(A1) or CA1-SR (B1) hippocampal region of 15 months old WT and Tg2576 treated with either vehicle (H_2_O) or PMX205. Scale bar: 5 μm. Quantitative analysis of VGlut1, PSD95 and colocalized synaptic puncta in the CA3-SL (A2-A4) or CA1-SR (B2-B4). Data are shown as Mean ± SEM (normalized to WT control group) of 3-4 images per mouse and n=4-5 mice per group. **C**. Imaris 3D reconstruction with surface rendering of individual microglial cells (Iba1+; grey), presynapses (VGlut1+; cyan), and lysosomes (CD68+; red). Scale bar: C1-C4: 5 μm; inserts 2 μm (C1-C4). Quantitative analysis of VGlut1+ presynaptic puncta engulfment per microglia (C5). Quantitative analysis of VGlut1+ presynaptic puncta localized within the microglial lysosomes (C6). Data are shown as Mean ± SEM of 15 individual microglial cells/mouse and n=4-5 mice per group. *p<0.05, ** p<0.01, *** p<0.001, **** p<0.0001.

We then investigated synaptic changes in the CA1-SR region in the Tg2576 mice treated with PMX205 and our results showed again a significant decrease of VGlut1 presynaptic puncta (22% decrease) in the Tg2576-H2O mice when compared to WT-H2O littermates (**Fig 5B2**). Treatment with PMX205 showed a trend towards the rescue of the excessive VGlut1 synaptic loss (Tg2576-PMX205 vs Tg2576-H_2_O; p = 0.14) and a significant increase in the PSD95 puncta in the Tg2576-PMX205 when compared to Tg2576-H_2_O (**Fig 5B3**). Colocalized VGlut1-PSD95 puncta showed a significant decrease (37%) in the Tg2576-H_2_O (vs WT-H_2_O) that (similar to the CA3-SL region) was rescued with PMX205 treatment (**Fig 5B4**). Interestingly, when single microglial cells in the CA1-SR region from all four groups were imaged and co-localization of VGlut1 within the microglial cell body or within the microglial lysosomes were quantified, a significant decrease in the amount of VGlut1+ presynaptic puncta within the microglial cell body in the Tg2576-H2O mice (compared to WT-H2O), suggesting that the loss of VGlut1 synapses observed in the CA1-SR region of the Tg2576 mouse model is not due to an excessive microglial synaptic pruning in this hippocampal region at this age (**Fig 5C5**). However, Tg2576-PMX205 treated mice did show reduced VGlut1 puncta localized within the microglial lysosomes (vs Tg2576-H_2_O mice) (Fig 5C6), consistent with our previous data in the Arctic model of AD and suggesting an improved lysosomal efficiency when C5aR1 signaling is suppressed. Overall, our results here demonstrated that pharmacological inhibition of C5a-C5aR1 signaling, by using a C5aR1 antagonist, can rescue the excessive synaptic loss observed in the Tg2576 mouse model of Alzheimer’s disease.

### 3.6 Astrocytes contribute to excessive pre-synaptic loss only at late stages of the disease

It is well established that astrocytes as well as microglial cells can participate in synaptic pruning in the healthy and disease brain, playing a key role in the pruning of synapses and circuit remodeling (23, 39, 45). Therefore, we assessed if C5a-C5aR1 signaling could influence synaptic engulfment by astrocytes, applying our two different approaches, the genetic deletion of C5aR1 in the Arctic mouse model of Alzheimer’s Disease (**Supp Fig 1A**) as well as the pharmacological inhibition of C5aR1 in the Tg2576 model of AD (**Supp Fig 1B**), where microglial ingestion of VGlut1 was not suppressed in CA1, while total VGlut1-PSD95 puncta was protected (**Fig 5B-C**). First, VGlut1 was co-stained with the astrocyte marker GFAP and the lysosomal marker Lamp2 and single astrocytes (and thus not plaque associated) were imaged at 10 and 7 months of age in the CA3-SL and the CA1-SR hippocampal region of WT, C5aR1KO, Arc and Arc-C5aR1KO mice. At 10 months of age in the CA3-SL region, while no difference in the VGlut1 puncta colocalized with GFAP+ astrocytic cell body in the Arctic mice vs the WT, Arc-C5aR1KO mice showed a significant reduction in the VGlut1 synaptic engulfment by astrocytes in the same age and region (vs Arctic) (**Fig 6A1**). Importantly, colocalization of VGlut1 puncta within the astroglial lysosomes showed a significant increase in the Arc mice when compared to WT littermates, that was reduced back to WT levels by the absence of C5aR1 (Arc-C5aR1KO vs Arc) (**Fig 6A2**), suggesting that, at least in the CA3-SL hippocampal region at this specific age, astrocytic synaptic pruning seems to be contributing to the excessive synaptic pruning, and that the absence of C5aR1 signaling suppresses this astrocyte engulfment of VGlut1. Similarly, at the same age (10 months) in the CA1-SR region, a significant reduction in the VGlut1 puncta located in the astroglial cell body was observed in the Arc-C5aR1KO vs Arc mice (**Fig 6A3**). Moreover, a significant increase in VGlut1+ puncta within the astroglial lysosomes (Arctic vs WT) and a trend for a reduction in the engulfment when C5aR1 was ablated (Arc-C5aR1KO vs Arc) was observed in the CA1-SR hippocampal region (**Fig 6A4**), which again, correlates with the excessive VGlut1 loss observed in the Arctic mice (**Fig 3A2 and C1)**). We then performed the same staining and quantification at 7 months of age in the same genotypes and our results showed a significant reduction of VGlut1 puncta colocalized with either the astrocyte cell body or the astroglial lysosomes in Arctic mice (when compared to WT controls) in the CA3-SL region (**Fig 6B1-B2**). In the CA1-SR area, we observed a significant reduction of VGlut1-GFAP colocalization in the Arc model (vs WT) (**Fig 6B3**), but no change in astrocyte lysosomal VGlut1. These results suggest that in contrast to microglia (**Fig 4C5**), astrocytes are not actively involved in excessive synaptic pruning and presynaptic loss observed in the Arctic mice at 7 months of age (**Fig 4**) and that C5aR1 deletion does not have an effect on astroglial synaptic engulfment at this stage in the Arctic model.

**Figure 6:**
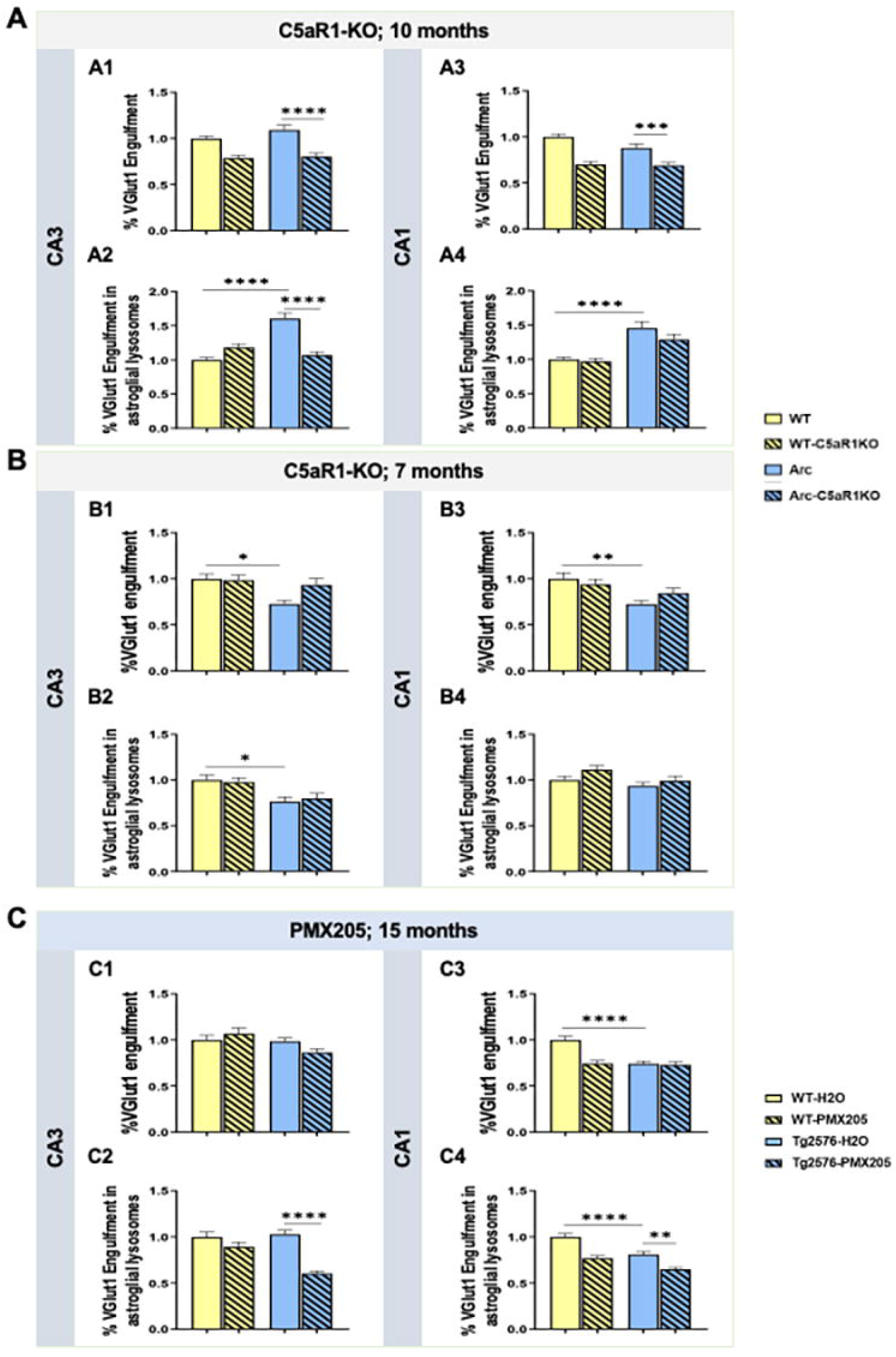
Contribution to VGlut1 synaptic engulfment by astrocytes differs based on age, region and the AD mouse model analyzed. **A.** Quantitative analysis of VGlut1+ presynaptic puncta engulfed per astrocyte (A1, A3) or localized within the astroglial lysosomes (A2, A4) at 10 months of age in WT, WT-C5aR1KO, Arc and Arc-C5aR1KO mice. Data are shown as Mean ± SEM (normalized to WT control group) of 15 individual astroglial cells/mouse and n=4 mice per genotype. **B**. Quantitative analysis of VGlut1+ presynaptic puncta engulfed per astrocyte (B1, B3) or VGlut1+ presynaptic puncta localized within the astroglial lysosomes (B2, B4) at 7 months of age in WT, WT-C5aR1KO, Arc and Arc-C5aR1KO mice. Data are shown as Mean ± SEM (normalized to WT control group) of 15 individual astroglial cells/mouse and n=3-4 mice per genotype. **C**. Quantitative analysis of VGlut1+ presynaptic puncta engulfed per astrocyte (C1, C3) or VGlut1+ presynaptic puncta localized within the astroglial lysosomes at 15 months of age in WT-H_2_O, WT-PMX205, Tg2576-H_2_O and Tg2576-PMX205 mice (C2, C4). Data are shown as Mean ± SEM (normalized to WT-H_2_O control group) of 15 individual astroglial cells/mouse and n=4-5 mice per genotype. *p<0.05, ** p<0.01, *** p<0.001, **** p<0.0001.

In contrast, the pharmacological inhibition of C5aR1 in Tg2576 and WT mice from 12-15 months of age, revealed no significant differences in the VGlut1 puncta localized in the astrocyte cell body in any of the experimental groups in the CA3-SL region (**Fig 6C1**), while a significant reduction in the lysosomal VGlut1+ puncta was observed in the Tg2576-PMX205 mice when compared to the Tg2576-H_2_O control group (**Fig 6C2**), suggesting that lack of C5aR1 signaling might enable increased lysosomal degradation activity. In the CA1-SR, we observed a reduction in VGlut1 engulfment by astrocytes (Tg2576-H_2_O vs WT-H_2_O) as well as the VGlut1+ puncta located in the astroglial lysosomes, which was further reduced by treatment with PMX205 (Tg2576-PMX205 vs Tg2576-H_2_O), similar to our observations in the CA3-SL region. These results suggest that astrocyte lysosomal activity is influenced as a result of C5a-C5aR1 signaling, but that astrocytes have a very limited role in synaptic elimination in the less aggressive amyloidogenic Tg2576 mouse models. Astrocyte contribution to the excessive synaptic pruning and synaptic loss were only seen at late stages (10 mo of age) of the disease in the aggressive Arctic48 mouse model of Alzheimer’s Disease.

## 4. Discussion

Synaptic loss is the strongest correlate for cognitive impairment in AD (18, 46, 47) and recent GWAS studies uncovered multiple AD risk variants that converge on phagocytic pathways in microglial cells (48). It is well-described that excessive complement activation can cause excessive or aberrant synapse pruning in mouse models of AD and other neurodegenerative disorders (16, 23, 49). Recent work has demonstrated that deletion of C1q, C3 and C4 can in fact, reduce the excessive synaptic loss described in multiple neurodegenerative disorders (19, 27, 28). However, the deletion of upstream components of the complement cascade also suppresses the generation of the downstream complement activation fragment C5a, and thus does not distinguish the contribution of C5a-C5aR1 signaling from the direct effects of the upstream components on synaptic pruning. In the present work we have investigated the role of C5a-C5aR1 interaction on synaptic pruning by using the Arctic48 mouse model of AD in which C5aR1 was genetically ablated, as well as the Tg2576 mouse model of AD upon treatment with a C5aR1 antagonist (PMX205). Here, in the more aggressive Arc model of AD there is an age and region-specific contribution of C5aR1 on synaptic pruning, as the deletion of this receptor results in reduced synaptic engulfment by microglial cells and a partial restoration of synaptic density in the CA3-SL region at 7 and 10 months of age, even without a change in C1q tagged synapses. Moreover, pharmacological inhibition of C5aR1 in the Tg2576 model of AD also rescued the excessive synaptic loss observed in this mouse model at 15 months of age in both CA3 and CA1. Thus, while C1q, complement activation and CR3 are critical in excessive synaptic pruning, these results indicate that the presence of C1q on the synapses is not necessarily sufficient to promote microglial engulfment, and that perhaps C5a-C5aR1 signaling induces changes in microglia engulfment capacity that is an important (and targetable) factor in excessive synapse loss.

These results are consistent with both the improvement in LTP that was observed here in the Arc-C5aR1KO mice, as well as the cognitive protection observed previously with C5aR1 antagonist treatment and the C5aR1 deficient Arctic model of AD (31, 33).

Synaptic pruning has been shown to occur by microglial engulfment through the CR3 receptor (20); however, the exact mechanism is still unclear. In fact, a recent work indicates that trogocytosis of presynaptic elements in a CR3-independent manner may also be involved (50). We have observed an age- and region-specific substantial loss of synapses in the Arctic and Tg2576 model of AD, that were largely due to decreases in the presynaptic markers SP or VGlut1. This decrease in presynaptic density correlated with higher VGlut1 engulfment by microglial cells. Genetic ablation or pharmacological inhibition of C5aR1 partially rescues the excessive microglial synaptic pruning and restores synaptic density in the CA3-SL, suggesting that microglial phagocytic capacity for presynaptic puncta could be a main driver involved in the synaptic density changes observed here. Synaptic pathology in Alzheimer’s disease has been reported to occur due to alterations in both the pre-synaptic and post-synaptic terminals (51); however, some studies, draw attention to earlier deficits at the pre-synaptic level in AD patients (43, 52, 53). In fact, a meta-analysis covering more than 400 publications (53) as well as an in-depth proteomics study pointed to a presynaptic failure in Alzheimer’s Disease patients (54).

In addition to microglial cells, it is well known that astrocytes have a vital role in synapse formation and maintenance and that they can also eliminate synapses during development but also in adulthood (38, 45, 55). Lee and colleagues demonstrated a major role for astrocytic Megf10 in eliminating excitatory synapses in the CA1 hippocampal region in adult mice (38). Moreover, it was recently proven that astrocytes preferentially eliminate excitatory synapses in a C1q-dependent manner in the Tau P301S tauopathy mouse model and that astrocytes can compensate for microglial dysfunction by increasing their engulfment of inhibitory synapses around amyloid plaques in the TauPS2APP;Trem2^KO^ mouse model (39). However, the specific role of astrocytes in synaptic pruning and synaptic loss during AD is still controversial and remains to be fully elucidated. Here we demonstrated a role for astrocytes in excitatory (VGlut1+) synaptic engulfment at late stages of the pathology in the Arctic48 model of AD, and more importantly, we showed that genetic ablation of C5aR1 reduces synaptic pruning by astrocytes. However, we did not observe a significant contribution from astrocytes in the excessive presynaptic pruning at earlier stages of the disease in the Arc mouse model or in the less aggressive Tg2576 model of Alzheimer’s Disease. Whether this is due to astrocytes not being activated enough at earlier stages of the pathology with limited amyloid deposition or to insufficient signaling from microglia to astrocytes to turn on their phagocytic mechanism remains to be determined.

C1q increases with age and disease in several mouse models and in AD (10, 42, 56). We have previously shown that C1q is locally synthesized by microglia and significantly increased in the Arctic model of AD (41). In addition, it is described that C1q tags synapses in several AD and tauopathy mouse models (19, 57) and that it preferentially tags the presynaptic side of the synapse, which can be directly linked to apoptotic changes in the synapse (58). In line with this, here we showed a significant increase in C1q tagging of presynapses at 7 and 10 months of age in Arctic mice relative to WT littermates. It has been hypothesized that C1q and iC3b tagged synapses are phagocytosed by microglia via CR3 receptor (20). Whether C1q associates with just “weak” synapses is not clear; however, stronger/more active synapses could be protected by complement regulatory proteins or by activity dependent factors (49, 59). We have observed here that the amount of C1q bound to synapses did not necessarily correlate with the changes observed in microglial synaptic engulfment or synaptic density due to C5aR1 deletion in the Arctic model of AD, supporting our hypothesis that the presence of C1q is not necessarily sufficient to induce synaptic ingestion. Additionally, we did not observe changes in C3 presynaptic deposition, which could suggest that the synaptic pruning requires additional signals (from C5a dependent microglial activation) or is counter-balanced by other protective signals. Others have previously presented data that there could be not only complement-dependent but also complement-independent mediated synapse pruning processes (16), which is also a logical extension from humans and models genetically lacking C1q or C3. Similar to our results here, Stephan and colleagues showed significant increases in C1q in adult mice with no synaptic loss associated with it. However, they did observe a circuitry reorganization, suggesting that C1q may also act independently of the complement cascade mediated synaptic elimination (42). It will be intriguing to determine if C1q at the high levels seen in AD would interfere with the several C1q-like molecules expressed in the nervous system that play roles in synapse formation and function (60, 61). In a very recent study from the Hong laboratory, they suggested a potential role of TREM2 upstream of C1q that could regulate microglial synaptic pruning in amyloidogenic models (62). Their results in pre-plaque stages of AD suggests that synapses externalized PtdSer in response to oAβ - induced injury and that microglia recognized PtdSer via TREM2 leading to removal of damaged synapses and reduced hyperactivity (62). This scenario fits to an extent with the findings of Zhong, et al, where TREM2 binding to C1q prevents complement activation and thus is protective in AD mouse models (63). However, further investigation is necessary to resolve confounding observations which may be related to the stage of disease in different mouse models as well as differences in levels of TREM2 function in these two reports. Whether TREM2 also plays a role in the excessive pre-synaptic elimination that we showed here or whether, and which, other “eat-me”,“don’t eat-me” and/or complement regulatory molecules (CD47, CSMD1/2, neuronal pentraxins) are involved are yet to be determined and will likely direct future therapeutic targets.

In summary, here we showed that the absence of C5aR1 (either by genetic ablation or by pharmacological inhibition) partially averts the excessive microglial synaptic pruning and the excessive presynaptic loss in two mouse models of Alzheimer’s disease, consistent with a significant role for C5a-C5aR1 signaling, downstream of early complement components, in these pathological events. We have previously shown that interaction of C5a with C5aR1 in microglial cells could produce a more inflammatory environment, suppressing beneficial clearance pathways that could contribute to neuronal damage and cognitive impairment (31). In addition, we have demonstrated a downregulation of a microglial cluster involved in synaptic pruning processes in Tg2576 mice treated with a C5aR1 antagonist (PMX205) (34), which could contribute to the results presented here. However, we cannot rule out the possibility that C5a-C5aR1 signaling could also impact neurons. C5aR1 expression has been shown in the stratum lucidum (64) where the mossy fibers coming from the granule cells of the DG contact pyramidal neurons of the hippocampal CA3 area. This pathway is involved in spatial memory formation and consolidation of hippocampal dependent memory (65). In fact, the presence of C5aR1 in the presynaptic terminals of the mossy fibers would enable a neuromodulatory role for C5a-C5aR1 signaling. Interestingly, it is precisely in the stratum lucidum of the CA3 region where we observed the restoration of VGlut1 pre-synaptic density when the C5a-C5aR1 axis is impaired. Regardless of the contributing mechanisms, modulation of C5a-C5aR1 signaling could both reduce the excessive synaptic pruning of “stressed-but-viable” (66) neuronal synapses and suppress neurotoxic inflammation, without suppressing the protective functions of the complement system and other innate immune systems both in the CNS and in the periphery. Importantly, avacopan, a small molecule C5aR1 antagonist (36, 67), has recently been granted FDA approval for a peripheral inflammatory disorder. Furthermore, PMX53, similar to PMX205, was shown to be safe in small phase 1b trial (68). Thus, precision targeting of C5aR1 is a promising novel therapeutic candidate for prevention and treatment of Alzheimer’s disease.

## Acknowledgements

Work was supported by NIH R01 AG060148, R21 AG061746 (AJT), R01 AG076835 (MAW) and U54 AG054349, Larry L. Hillblom postdoctoral fellowship #2021-A-020-FEL (AGA) and Alzheimer’s Association Research Fellowship AARFD-20-677771 (NDS). We thank Enrico Gratton and Rachel Cinco in the Laboratory for Fluorescence Dynamics, Department of Biomedical Engineering University of California, Irvine, CA for their help and use of the Airyscan especially in the early stages of this project. This study was made possible in part through access to the Optical Biology Core Facility of the Developmental Biology Center, a shared resource supported by the Cancer Center Support Grant (CA-62203) and Center for Complex Biological Systems Support Grant (GM-076516) at the University of California, Irvine.

## Disclosures

Authors declare no conflict of interest.

## Abbreviations

AD: Alzheimer’s disease
Arc: Arctic48
DG: dentate gyrus
LTP: long-term potentiation
ML: molecular layer
PSD95: postsynaptic density protein 95
SL: stratum lucidum
SP: synaptophysin
SR: stratum radiatum
WT: wild type

---

**Supplemental Figure 1: Experimental design.**

**A.** Schematic diagram of the experimental design showing that Arc48+/- mice were crossed with C5aR1-KO mice to create Arctic mice lacking C5aR1 (Arc-C5aR1KO). For the experiments shown in this manuscript, WT, WT-C5aR1KO, Arc and Arc-C5aR1KO mice were aged to 2.7, 5, 7 and 10 months of age. Amyloid pathology is prominent in this mouse model at 4 mo of age. **B**. Schematic diagram of the experimental design showing that Tg2576 mice and WT littermates were treated with 20 µg/ml of PMX205 in drinking water (or H_2_O) for 12 weeks from 12-15 months of age, the time corresponding to the onset and accumulation of amyloid pathology in this mouse model of AD.

**Supplemental Figure 2: Specificity of rabbit monoclonal anti-mouse C1q** Representative super-resolution images of VGlut1 (green), C1q (red) and merged C1q-VGlut1 in the CA1-SR hippocampal region of Arctic-C1qa^FL/FL^ (top panel) and C1qa deleted Arctic-C1qa^FL/FL^:Cx3cr1^ERT2^ (bottom panel) at 10 months of age. Scale bar: 5 μm.

**Supplemental Figure 3: Loss of neuronal complexity in CA3 is prevented by C5aR1 ablation in the Arctic model of AD at 10 months.**

Sholl analysis of WT, Arctic, C5aR1-KO, and Arctic C5aR1-KO in the CA3 area at 10 months of age. Neurite length intersections between 0 and 200 µm from the soma were averaged in 10-20 neurons per mouse from 3-4 mice per genotype. Data are shown as Mean ± SEM. *p<0.05 when comparing Arc vs Arc-C5aR1KO. In addition, using Kolmogorov-Smirnov test, WT vs Arctic, p=0.006; Arc vs Arc-C5aR1KO, p=0.002; whereas WT vs Arc-C5aR1KO is not significantly different.

**Supplemental Figure 4: C3 tagging of VGlut1 presynaptic puncta at 10 months of age.**

**A.** Representative super-resolution images (A1) and quantitative analysis of C3, VGlut1 and C3-VGlut1 colocalized puncta (A2-A4) in the CA3-SL region. **B**. Representative super-resolution images (B1) and quantitative analysis of C3, VGlut1 and C3-VGlut1 colocalized puncta (B2-B4) in the CA1-SR region. Scale bar: 5 um. Data are shown as Mean ± SEM (normalized to WT control group) of 3 images per animal and n=3-4 animals per genotype. *p<0.05; **p<0.01.

**Supplemental Figure 5: No differences in synaptic density despite an increase in C1q tagging of presynaptic puncta in the DG-ML at 10 months of age.**

**A.** Super-resolution images of SP (green) and PSD95 (red) in the DG-ML of 10m Arc, Arc-C5aR1KO and their respective WT littermates. Scale bar: 5 μm (A1). Imaris quantification of SP, PSD95 and SP-PSD95 co-localized puncta (A2-A4). Data are shown as Mean ± SEM (normalized to WT control group) of 3 images per animal and n=2-3 animals per genotype. **B**. Quantitative analysis of C1q, C1q-VGLUT1 and C1q-SP colocalized puncta. Data are shown as Mean ± SEM (normalized to WT control group) of 3 images per animal and n=1-4 (C1q-SP) or n=3-7 (C1q-VGLUT1) mice per group. * p<0.05; **p<0.01.

**Supplemental Figure 6: VGlut1 microglial engulfment or synaptic density remain unchanged despite an excessive C1q tagging in the CA1-SR hippocampal region at 7 months in the Arctic model of AD.**

Quantification of super-resolution images of (**A**) VGlut1, PSD95 and colocalized VGlut1-PSD95 puncta or (**B)** of VGlut1 and C1q-VGlut1 puncta in CA1-SR at 7 months of age. Data are shown as Mean ± SEM (normalized to WT control group) of 3 images per mouse and n=3-4 mice per genotype. **C**. Confocal images and 3D surface rendering of Iba1, CD68 and VGlut1 engulfment at 7 months of age in the CA1-SR region of the hippocampus. Scale bar: 10 μm; inserts 2 μm (C1-C4). Quantitative analysis of VGlut1+ presynaptic puncta engulfment per microglia (C5) or localized within the microglial lysosomes (C6). Data are shown as Mean ± SEM of 15 individual microglial cells/mouse and n=3-4 mice per genotype. *p<0.05, ** p<0.01, *** p<0.001, **** p<0.0001.

**Supplemental Figure 7: No differences in synaptic density were observed in CA3-SL or CA1-SR at 2.7 or 5 months of age.**

**A-B**. Quantitative analysis of colocalized VGlut1-PSD95 puncta at 2.7 months (A) or 5 months (B) of age in the CA3-SL (A1, B1) and CA1-SR (A2, B2) hippocampal regions. Data are shown as Mean ± SEM (normalized to WT control group) of 3 images per animal and n=3 animals per genotype.

## Notes

### Competing Interest Statement

The authors have declared no competing interest.

