## Supplementary figures and images for "C5aR1 signaling promotes region and age dependent synaptic pruning in models of Alzheimer’s Disease"

### Supplemental Figure 1

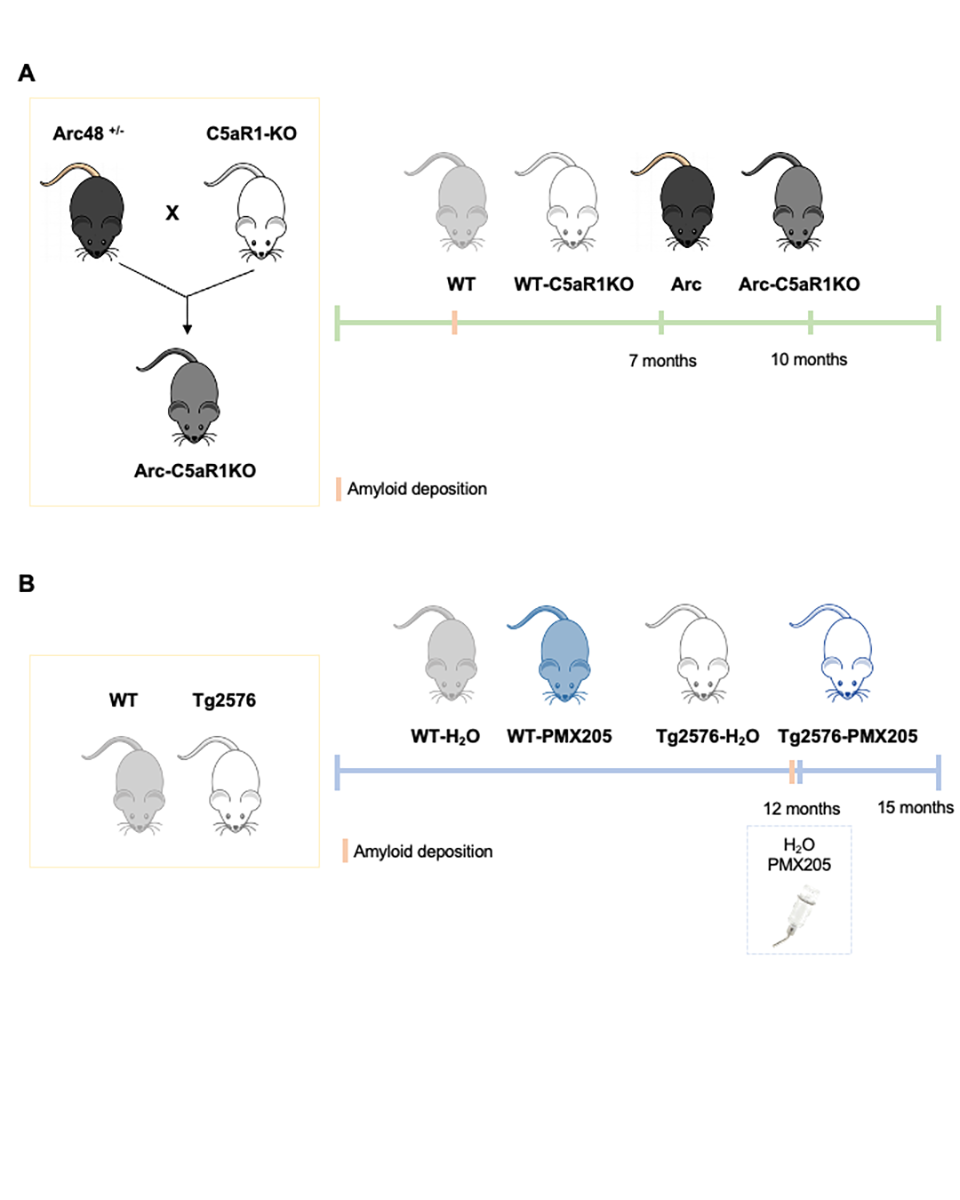

### Supplemental Figure 2

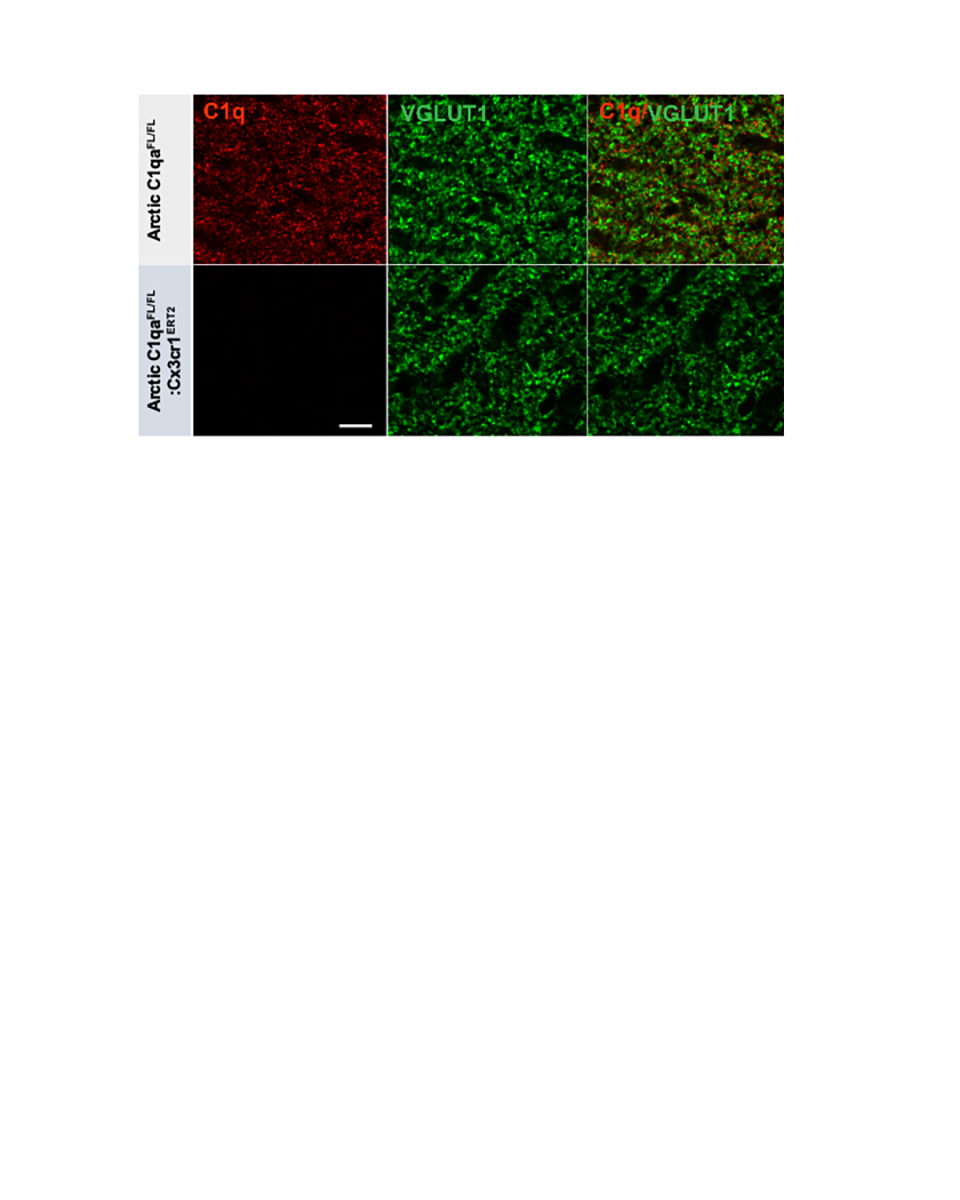

### Supplemental Figure 3

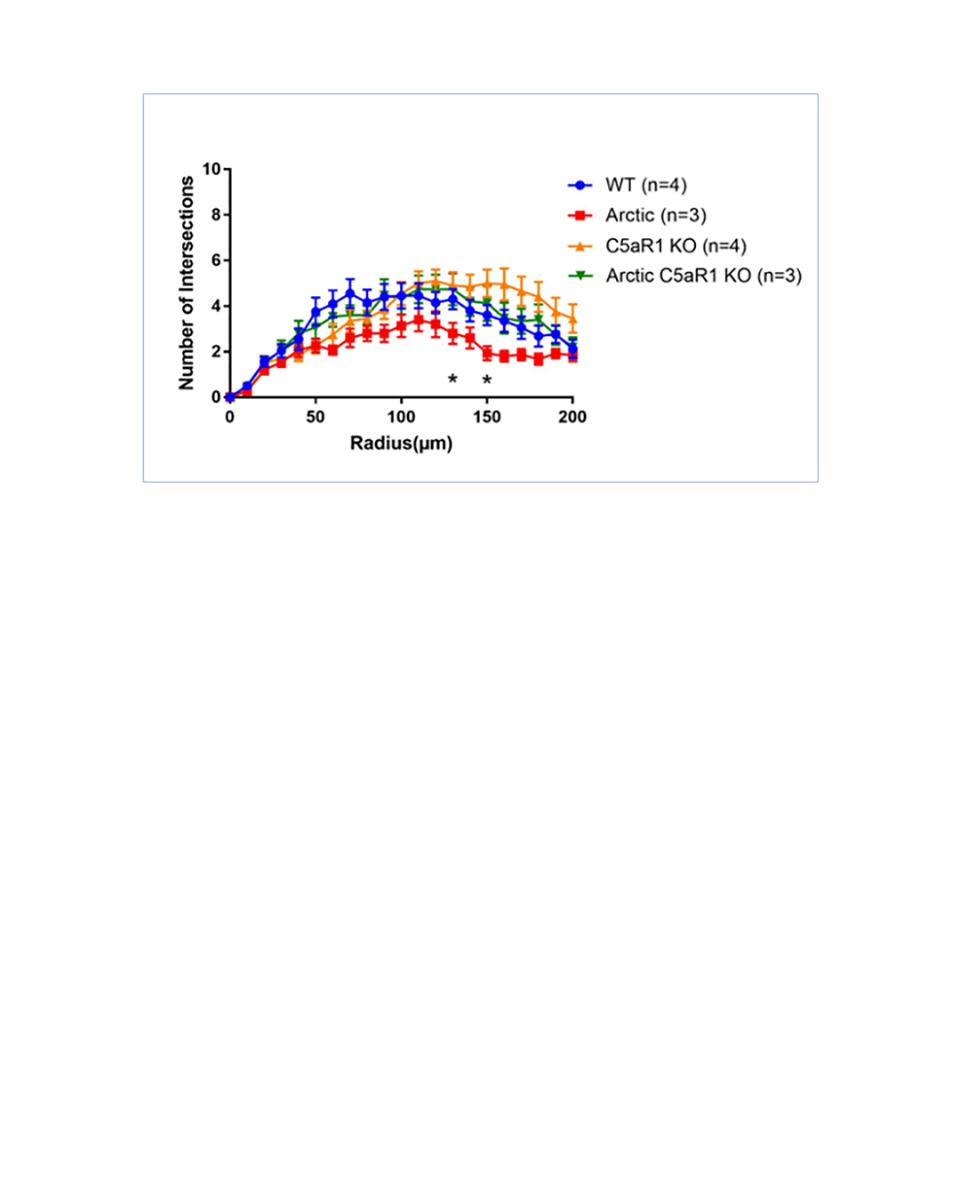

### Supplemental Figure 4

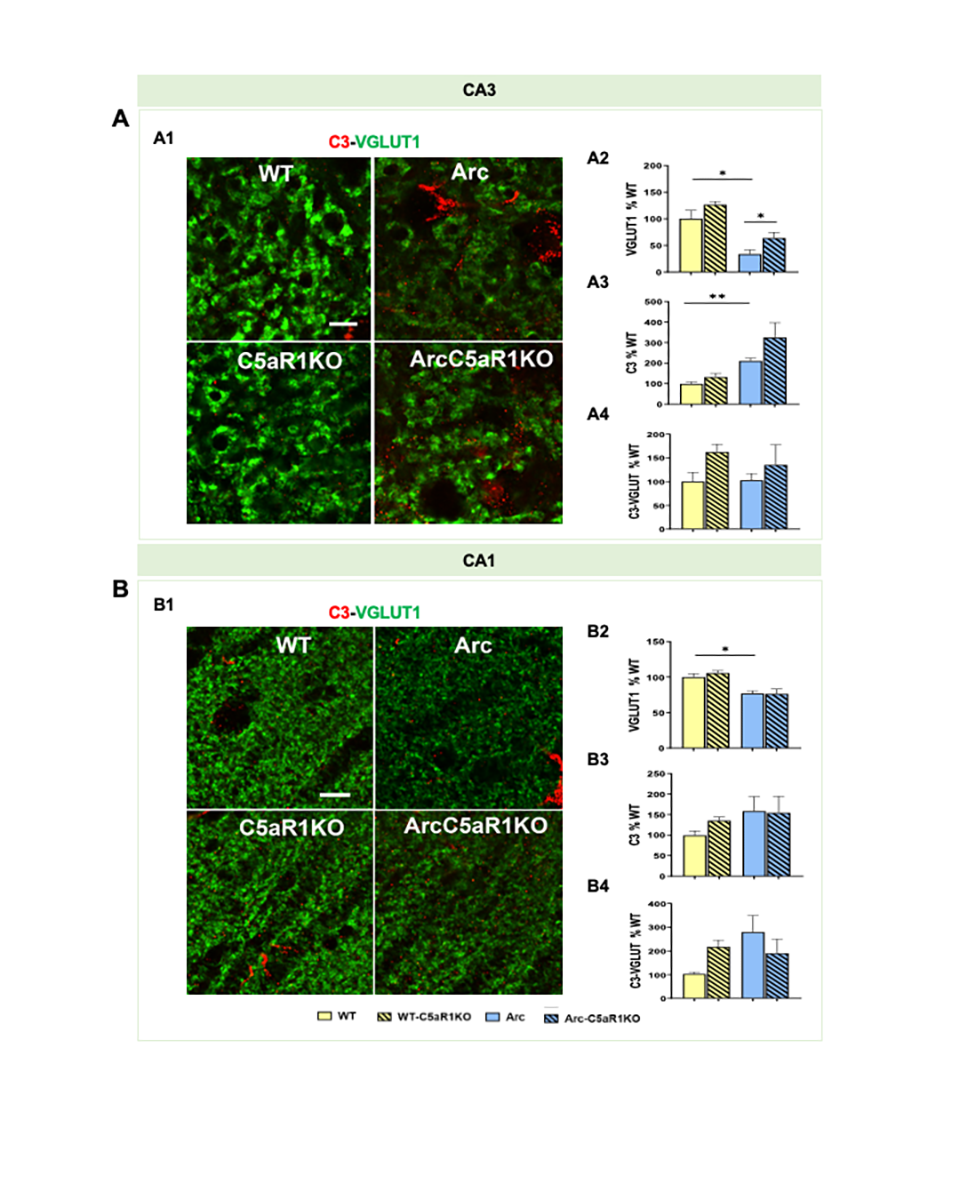

### Supplemental Figure 5

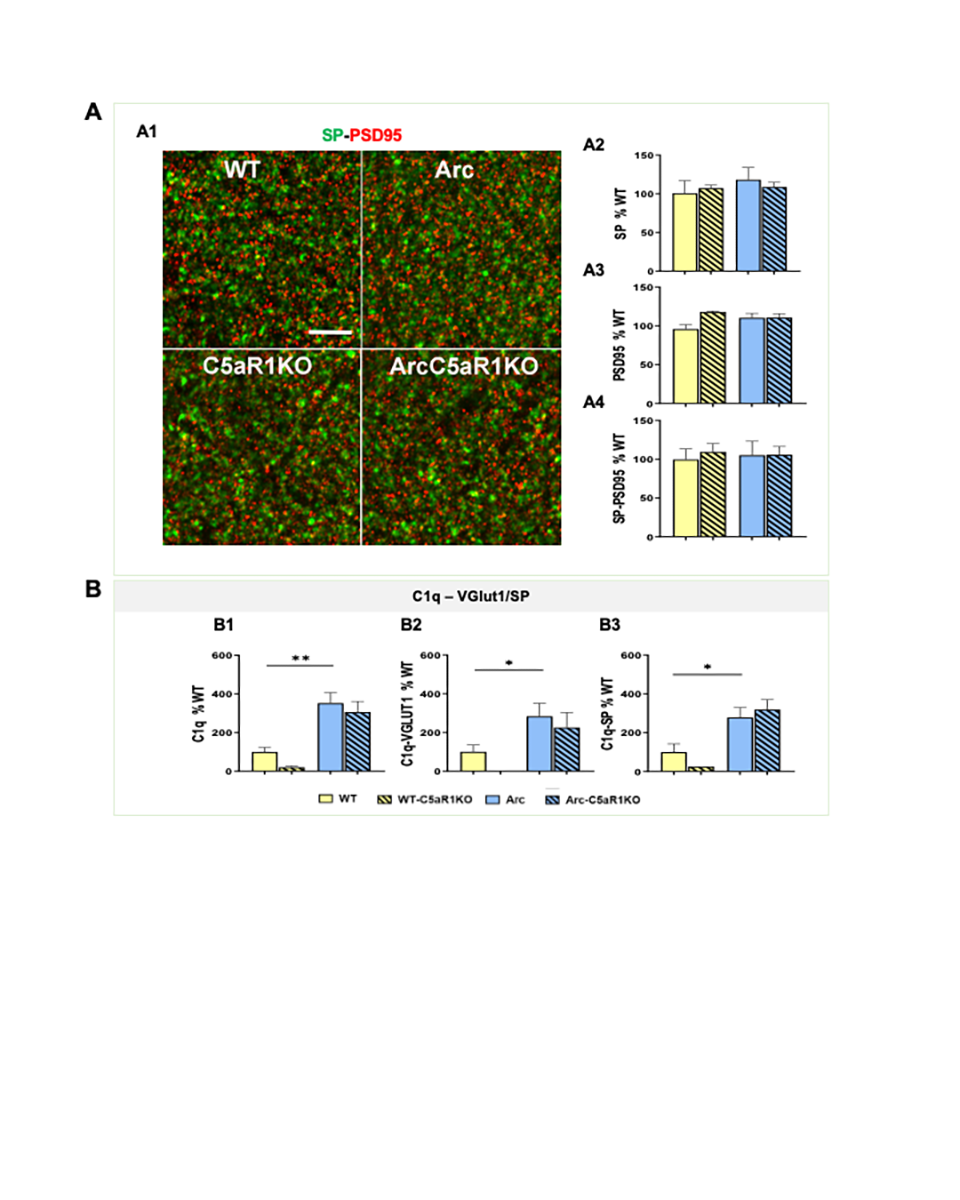

### Supplemental Figure 6

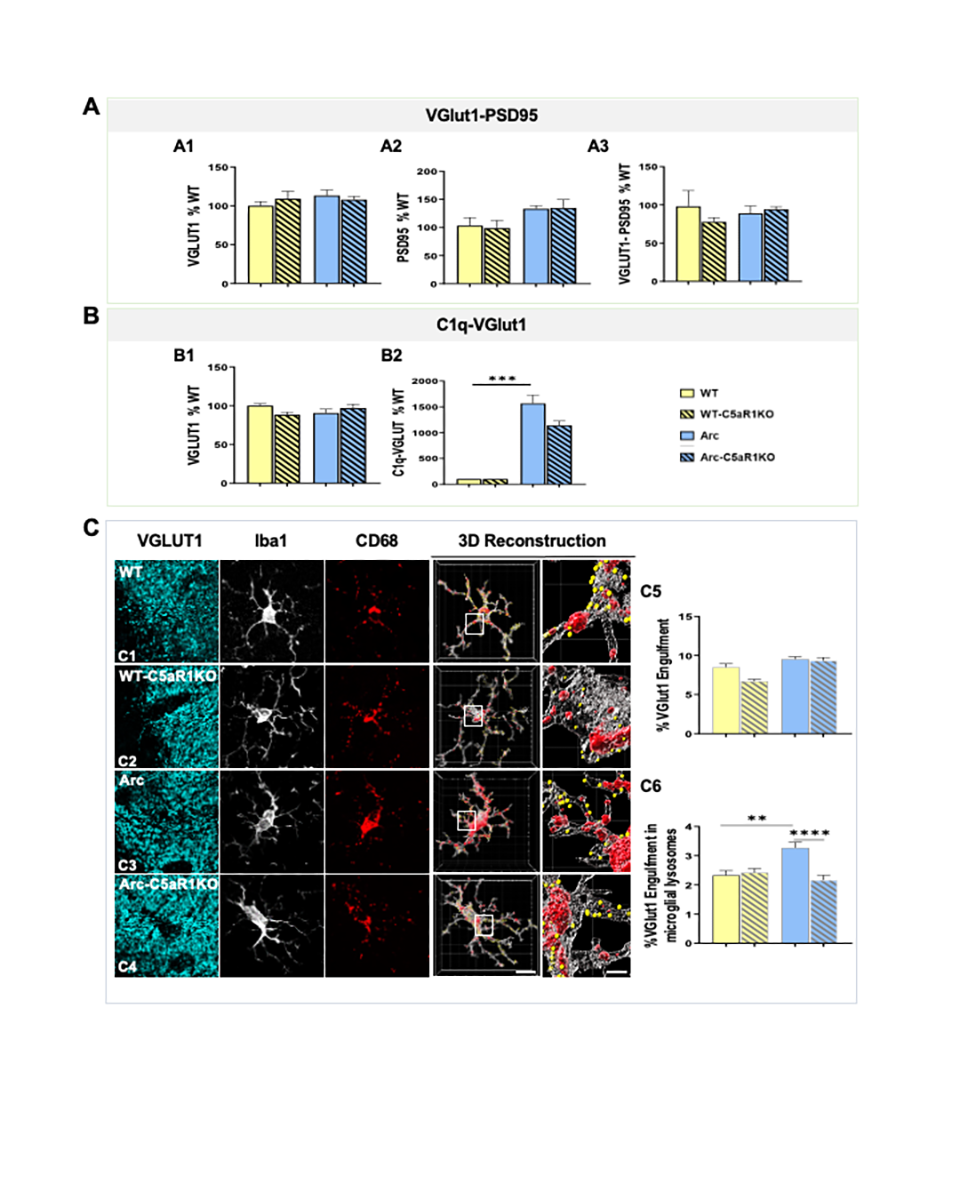

### Supplemental Figure 7

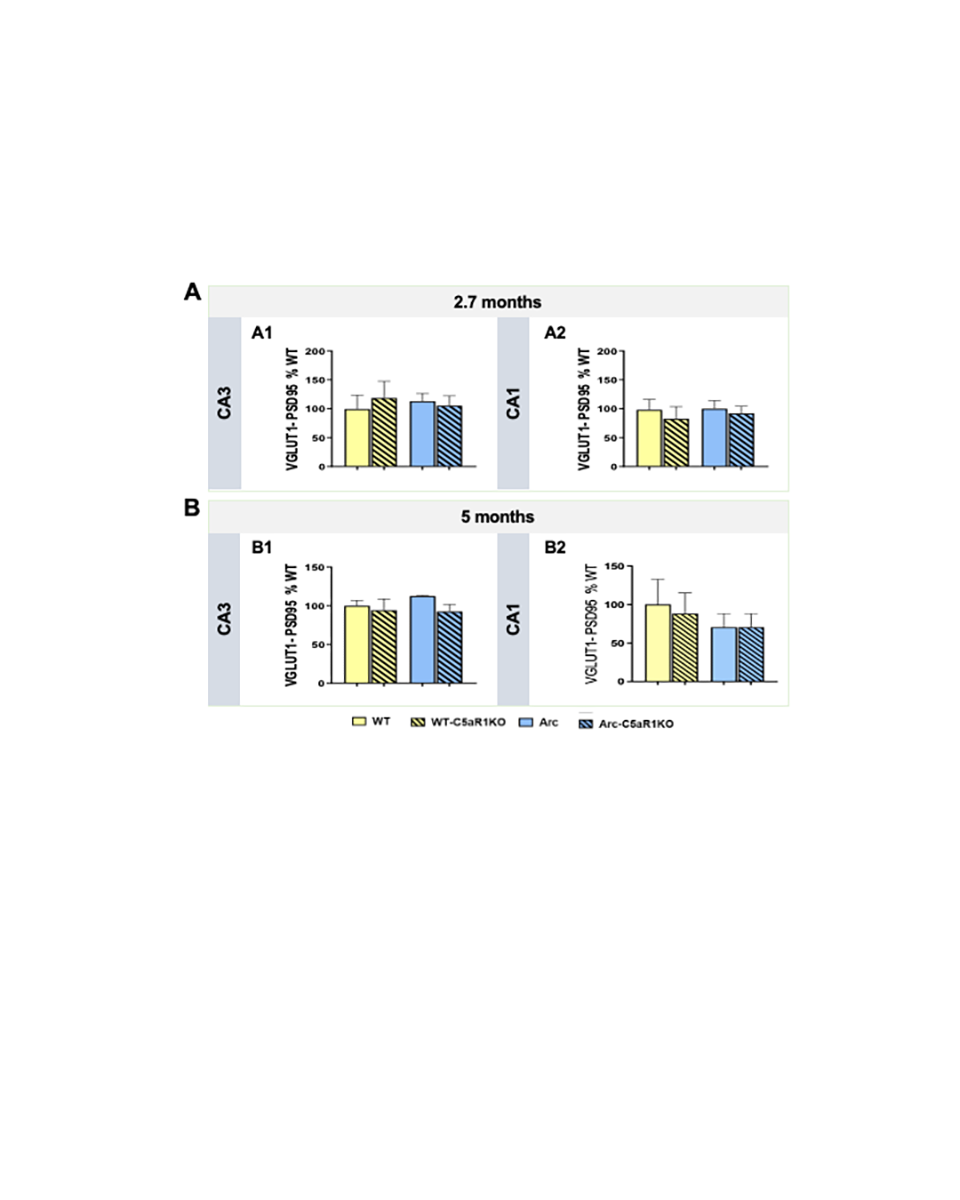
